# Uromodulin promotes immune zonation and inhibits alternative inflammasome-mediated activation of immune-to-collecting duct inflammatory signaling in early acute kidney injury

**DOI:** 10.64898/2026.02.26.708299

**Authors:** Angela R. Sabo, Azuma Nanamatsu, Dillen Wischmeier, Connor J. Gulbronson, Shehnaz Khan, Radmila Micanovic, Seth Winfree, Tarek El-Achkar, Kaice A. LaFavers

## Abstract

Uromodulin, a protein made uniquely by the kidney, is protective against acute kidney injury. An integrated transcriptomic and multiplexed spatial protein imaging was used to uncover early cellular and molecular pathophysiological mechanisms following murine kidney ischemia and reperfusion injury (IRI) and better define the role of Uromodulin at the early stage of injury. Six hours following IRI, there was a pan-nephronal transcriptomic response with activation of common pathways but also unique gene expression signatures for each nephron segment. Cell-cell communications and epithelial-immune spatial interactions most prominently involved thick ascending limbs and distal nephron segments with distinct immune zonation in the inner stripe of the outer medulla. Uromodulin deficiency swayed the tubular transcriptomic signatures towards more severe injury and inflammation with altered macrophage communication. Uromodulin deficiency also caused partial loss of immune zonation and a shift towards broader epithelial-immune interactions in the outer stripe and cortex. Uromodulin inhibited activation of the *Nlrc4-*dependent alternative inflammasome pathway in macrophages, where the production of IL-1β predominantly targets other immune and collecting duct (CD) cells. Indeed, Uromodulin deficiency induced the expression of CD8 in CD cells which acquire a proinflammatory phenotype linked to spatial niches containing immune cells. The presence of CD8^+^ CD cells was validated in human kidney biopsies. In conclusion, our findings support a role for Uromodulin in spatially confining the immune system around TAL cells in the inner stripe away from the vulnerable outer stripe in early injury. Uromodulin also inhibits the inflammasome-mediated macrophage-epithelial crosstalk that could induce collecting duct cells towards more inflammatory signaling.

## Introduction

Acute kidney injury (AKI) is a serious disease frequently affecting hospitalized patients^1^. AKI has grave implications for long term kidney disease development and progression to chronic kidney disease (CKD)^2^. Newer technical approaches such as single cell RNA sequencing and spatial transcriptomics have been recently used to characterize the transition from AKI to CKD in experimental models of kidney injury and using human kidney biopsy specimens of patients with kidney disease. This has led to characterization of various states of repair in epithelial cells, including defining states of failed repair associated with ongoing activation of the immune system leading to disease progression and fibrosis^3–7^. Important insights using various experimental models of AKI such as ischemia-reperfusion or nephrotoxic injury have also uncovered key aspects of the pathogenesis of AKI, especially at the peak of injury^8^. These studies suggest that early changes shortly after the onset of injury could be key determinants governing the severity of injury, which in turn could determine the course of recovery. However, investigations using novel omics technologies, particularly spatial protein imaging, have not been used comprehensively to study these early cellular events *in situ*. Understanding such events could also lead to identifying factors associated with increased risk of AKI for prevention purposes.

Uromodulin (UMOD, also known as Tamm-Horsfall Protein) is a glycoprotein uniquely made in the kidney by cells of the thick ascending limbs (TAL) of the loop of Henle and early distal convoluted tubules^9,10^. From a cell biology perspective, UMOD is predominantly secreted from the apical domain in the urinary filtrate as the most abundant protein in healthy urine^11^. Urinary UMOD is present mostly in a polymerized form and is thought to play a key role in protection from urinary tract infections and stone disease^12,13^. However, UMOD is also released to a lesser extent from the basolateral domain towards the renal interstitium and circulation, where it exerts immunomodulatory properties and regulates systemic oxidative stress^14–16^. Indeed, circulating UMOD, which is predominantly monomeric, inhibits inflammatory signaling in proximal tubules and promotes macrophage activation^16,17^. UMOD deficiency is associated with increased AKI susceptibility and severity^18^. Therefore, studying early spatial and molecular changes after acute injury during UMOD deficiency will also uncover key pathways that sway kidney injury towards a more severe phenotype and enhance our understanding of the immunomodulatory role of UMOD.

Spatial molecular imaging technologies have become more widely available, yielding important advancements in our understanding of the course of AKI. Unbiased and targeted spatial transcriptomics have given insights into the association of tubular cells that are failing to repair with foci of inflammation or fibrosis^6,19,20^. However, immune cells are detected with variable frequency by such methods. To circumvent that, the signature of immune cells from single cell RNA sequencing is commonly transferred *in situ*, to localize with better certainty the distribution of immune cells^21^. Spatial protein imaging such as co-detection by indexing (CODEX) or the newer Phenocycler-Fusion platform are increasingly used to image protein at the sub-cellular level. They provide the advantage of faithfully detecting immune cells with high resolution^22^ and incorporating markers that reflect true active state such as phospho-proteins and nuclear localization of transcription factors. We recently used spatial protein imaging in human kidney biopsies to track the changes of proximal tubules expressing stem cell markers in kidney disease and link their spatial association with immune cells^23^. Neighborhood analysis using this technology can uncover important microenvironments that can elucidate the biology and patho-mechanisms of disease, especially when linked to ligand-receptor analysis from single cell or spatial transcriptomics^24^. Spatial protein imaging has not been applied at this scale in experimental kidney disease particularly in early stages post injury.

In this work, we integrated spatial protein imaging with CODEX and single cell RNA sequencing to define immune spatial signatures and niches that characterize health and study early events post kidney injury. We uncover unexpected immune zonation and involvement of the distal nephron, which exposes the complexity of the coordinated response to protect the most vulnerable parts of the kidney after injury. We also define key changes occurring with UMOD deficiency, whereby inflammasome-mediated immune-epithelial crosstalk triggers a potent inflammatory response in specialized collecting duct cells.

## Materials and Methods

### Study Design

In this study, we studied the early response to ischemia reperfusion in UMOD^-/-^ mice to generate, and subsequently test hypotheses about UMOD’s protective role in acute kidney injury. We performed bilateral ischemia reperfusion injury (IRI) surgeries on UMOD^+/+^ and UMOD^-/-^ mice and isolated samples for single cell transcriptomics and multiplexed imaging for hypothesis generation. Additional experiments were performed to validate, explore and expand on these results, as described in the following methods sections. Sample sizes were established for different analyses as follows. For *in vitro* experiments, all experiments were performed with three technical replicates and were repeated a minimum of 1-2 additional times to ensure reproducibility. For animal experiments using the IRI model of AKI, experiments were conducted with 3-5 mice per group, based on the number of mice required for significance in uninjured animals along with feasibility considerations. The subjects were not randomly assigned to experimental groups due to animal housing considerations. The study was not blinded. All data acquired is presented in the main figures and supplementary data. These criteria were established prospectively. Statistical outliers were defined using the Tukey’s fences methods, unless otherwise described in the specific methods sections.

### Mice

Animal experiments and protocols were approved by the Indianapolis VA Animal Care and Use Committee. Age matched 8-12 week-old Uromodulin (UMOD) knockout male mice (129/SvEv UMOD^-/-^) and wild type background strain (129/SvEv UMOD^+/+^) were used as described previously^14^. Ischemia reperfusion injury was performed as described previously^14^. Briefly, mice were anesthetized with inhaled isoflurane in an induction chamber. An abdominal incision was made to access the mouse kidneys, whose renal pedicles were then clamped for 22 or 30 minutes. For the sham surgery, mice experience identical surgical preparation, incision, and recovery without clamping. Mice were allowed to recover for six hours before euthanasia and tissue harvest.

### Measurement of Serum Creatinine

Concentrations of mouse serum creatinine were measured by isotope dilution liquid chromatography tandem mass spectrometry by the University of Alabama O’Brien Center Bioanalytical Core facility.

### Dissociation of Kidney into Single Cells

The protocol used for kidney dissociation was provided by Janosevic et al^25^. Briefly, both kidneys from one male 129Sv/Ev mouse were removed after a sham surgery or two males after ischemia surgery and minced together on a glass petri dish into ∼8 pieces total and placed in Miltenyi Biotec tubes (C-tubes) with tissue digestion mix from the Multi Tissue Dissociation Kit 2 (Miltenyi Biotec) and agitated by a gentleMACs Dissociator (Miltenyi Biotec) followed by rotation on the MACSMix Tube Rotator (Miltenyi Biotec) for 30 minutes at 37°C and a final agitation cycle on the gentleMACs Dissociator. RPMI with 5% BSA was added to quench the enzymatic reactions and the suspension was passed through a 40 μm strainer that was washed with 5 mL RPMI with 0.04% BSA. The eluent was passed through a second 30 μm strainer and wash with 5 mL of RPMI with 0.04% BSA. The total eluent was centrifuged at 300 x g for 5 minutes at 4°C and the pellet resuspended in 1 mL chilled RBC Lysis Buffer (Sigma) and incubated for 5 minutes at 4°C. 10 mL cold RPMI with 0.04% BSA was added to dilute the lysis buffer and the sample was spun at 1000 rpm in a tabletop centrifuge at 4°C (Eppendorf 5430). The cells were washed three times with 5-minute spins to pellet the cells. After the final wash the pellet was resuspended in 1 mL of cold RPMI with 0.04% BSA. Dead cells were removed from the suspension using the EasySep Dead Cell Removal by Annexin V Kit (Stemcell Technologies). The magnetic beads were isolated using a multi-tube magnet for 1.5 mL tubes (Invitrogen). After removal of the dead cells the cell suspensions were transferred to 15 mL conical tubes and centrifuged at 1200 rpm in the Eppendorf 5430 tabletop centrifuge at 4°C. The cell pellet was resuspended in 1-3 mL of RPMI with 0.04% before running on the Chromium 10X platform. processing.

### Kidney Dissociation and Isolation of CD45^+^ Kidney Cells

Kidneys were homogenized with a Tissuemiser (Fisher Scientific) handheld homogenizer on ice by making 10 passes three times. The tissue slurry was incubated with collagenase type IA for 30 min at 37°C. The resulting suspension was passed through a 70 µm strainer and washed with PBS + 1% fetal bovine serum (FBS) before pelleting at ∼1,600 x g for 10 min. To remove red blood cells and cellular debris the suspension was clarified on a Percoll gradient ^26^. At room temperature, the cell pellet was resuspended in 8 mL of 40% Percoll in PBS with 1% FBS and layered on top of 3 mL of 70% Percoll in PBS with 1% FBS at room temperature. The gradient was spun for 30 minutes at 900 x g at room temperature. The centrifuge brake was turned off to not disturb the formed gradient and separated cells. Following the spin there was a fatty cellular debris layer at the top of the gradient, a fuzzy “buffy coat” at the interface between the 40% and 70% Percoll, a thin band of red blood cells just below and a pellet of red blood cells. The buffy coat was removed and diluted to 15 mL with PBS with 1% FBS and spun at 800 x g for 10 min at 4°C. After removing the supernatant, the pellet was resuspended in 1 mL PBS with 1% FBS. In preparation for labeling with antibodies for sorting by the flow cytometry facility, the suspension was centrifuged and resuspended in 100 µL to which CD16/32 antibody (BDBioscience) was added and incubated for 5 minutes at room temperature to block Fc gamma receptors. To label the cells with CD45 antibody, 10 µL (1.5 µg) APC conjugated CD45 antibody (clone 30F11, Miltenyi) was added to the 100 µL suspension and incubated for 30 minutes at 4°C. Cells were resuspended in PBS with 1% FBS and mixed with an equal volume of 1 µg/mL propidium iodide. CD45^+^ live cells were selected on the basis of PI and APC CD45 staining. The sorted cells were collected on ice.

Dead cells were removed using the Easy Sep Dead Cell (Annexin V) Removal Kit, according to the manufacturer’s direction. To remove calcium salts, the cells were washed twice with PBS with 1% FBS and filtered (40 µm filter) before running on the Chromium 10X platform.

### Analysis of scRNA Seq Data

Sequencing data was analyzed using the Seurat package (Ver 5.2.1)^27^ in R Studio, with ligand-receptor analysis performed using the CellChat package (Ver 2.2.0)^28^ in R Studio. Raw and processed data are available through the Gene Expression Omnibus repository under the following datasets: GSE171639, additional samples to be added to a repository upon acceptance of this manuscript.

### CO-Detection by IndEXing – Immuno-fluorescence multiplexed imaging and image analysis

Mouse kidneys were cut in half and preserved in OCT compound at -80°C. Ten-micrometer thick sections were then cut onto poly-L-lysine coated coverslips and processed using the protocol provided by Akoya Biosciences and as described previously by Melo-Ferreira *et al*^20^. In brief, sections were rehydrated with a three-step hydration process, followed by multiple fixation steps, overnight staining with an antibody cocktail (**Table 1**), and post-staining washes and fixes. Oligonucleotide probe staining and fluidics were handled using the CODEX 1.0 system from Akoya Biosciences and imaging was performed using a Keyence BZ-X810 automated tile-scanning microscope. The resulting images were processed using the CODEX processor from Akoya Biosciences and visualized using FIJI/ImageJ. After processing, images were segmented based on DAPI signal using Volumetric Tissue Exploration and Analysis (VTEA)^29^. Initial semi-unsupervised clustering was done in VTEA to remove artifacts from analysis such as tissue folds or “cells” that were positive for too many markers. After samples were “cleaned”, combined clustering and neighborhood analysis was done in R using a variety of packages indicated in the code provided (to be deposited in Zenodo upon acceptance of manuscript).

**Table 1.** Antibodies in multiplex panel.

| **Protein Target**<br>**(Mouse)** | **Manufacturer** | **Antibody Clone** | **Research Resource**<br>**Identifier** |
| --- | --- | --- | --- |
| Nephrin | Novus Bio | polyclonal | AB_2155023 |
| VWF | Novus Bio | polyclonal | AB_10002749 |
| Endomucin | Abcam | V.7C7.1 | AB_10859306 |
| αSMA | Invitrogen | 1A4 | AB_2574460 |
| CD31 | Akoya Biosciences | MEC13.3 | AB_3254603 |
| LRP2 | Abcam | ab76969 | AB_10673466 |
| UMOD | R and D Systems | polyclonal | AB_2212556 |
| Na<sup>+</sup>/K<sup>+</sup> ATPase | GeneTex | HL114 | AB_2888530 |
| AQP2 | ThermoFisher Scientific | PA5-22865 | AB_11154667 |
| CD3 | Akoya Biosciences | 17A2 | AB_3739919 |
| CD4 | Akoya Biosciences | RM4-5 | AB_3271537 |
| CD8a | Akoya Biosciences | 53-6.7 | AB_3271540 |
| CD11b | Akoya Biosciences | M1/70 | AB_3271542 |
| CD11c | Akoya Biosciences | N418 | AB_3739920 |
| CD74 | R and D Systems | AF7478 | AB_2941942 |
| CD169 | Akoya Biosciences | 3D6.112 | AB_3739921 |
| CD206 | BioLegend | CO68C2 | AB_10900263 |
| LY6G | Akoya Biosciences | 1A8 | AB_3739925 |
| MHCII | Akoya Biosciences | M5/114.15.2 | AB_3271532 |
| CD45 | Akoya Biosciences | 30-F11 | AB_3271536 |
| NKp46 | BioLegend | 29A1.4 | AB_10551441 |
| B220 | Akoya Biosciences | RA3-6B2 | AB_3739928 |
| CD71 | Akoya Biosciences | R17217 | AB_3739929 |
| Ki67 | Akoya Biosciences | B56 | AB_2895046 |
| KIM-1 | R and D Systems | AF1817 | AB_2116446 |

### Immuno-fluorescence confocal microscopy and 3D tissue cytometry

Immuno-fluorescence staining for LY6G (Biolegend, Clone 1A8), HAVCR1 (also known as Kidney Injury Molecule 1 or KIM-1, R&D Systems, Cat #MAB1750), ATF3 (Cell Signaling Technologies, Cat # 18665), DAPI (Sigma Aldrich, Cat #D9542), and FITC-phalloidin (Molecular Probes, Cat. #D1306 and #F432) was performed on 50 µm vibratome sections of kidneys fixed with 4% para-formaldehyde as described previously^14^. Briefly, immunostaining was done at room temperature, overnight, with primary antibody in PBS + 2% BSA + 0.1 % Triton X-100 and secondary antibody + FITC-phalloidin + DAPI in the same buffer. After mounting on glass slides, stained sections were viewed under Leica SP8 upright confocal microscope with a motorized stage, 2 HyD and 2 PMT detectors with 4 lasers at 405 nm, 488 nm, 552 nm and 635 nm controlled by LASX software. Automated tile scanning and mosaic imaging was performed in four channels to cover the whole tissue in x, y, and z. The captured volumes were stitched in LASX and 3D tissue cytometry was performed with Volumetric Tissue Exploration and Analysis (v0.5.2)^30^. Briefly, cells were segmented by their nuclei. Identifying cell-types or cell characteristics was accomplished by measuring the intensity of markers either within the putative nuclei or a small surrounding volume generated with a morphological dilation. These measurements are collated and plotted on an interactive scatter plot which can be interrogated with gating approaches. Here, HAVCR1/KIM-1 and ATF3 intensity were assessed in the nuclei of segmented cells and positive cells were gated and counted.

### Measurement of IL-1β

Levels of circulating IL-1β were measured by ELISA. Samples were diluted 1:4 and assayed according to the manufacturer’s protocol (R&D Systems, MLB00C-1).

### Preparation of Bone Marrow Derived Macrophages

Bone marrow derived macrophages were generated as described previously^31^. Briefly, femurs and tibia from euthanized UMOD^+/+^ mice were harvested aseptically into sterile PBS/1% FBS. Excess muscle was removed, and the leg bones were severed proximal to each joint. A 25-gauge needle was used to flush each bone with cold sterile PBS with 1% FBS. Cells were centrifuged for 10 min at 500 x g (room temperature). Supernatant was discarded and the cell pellet resuspended in RPMI 1640 with L-glutamine, 10% FBS and 1X pen/strep. Bone marrow cells were differentiated into macrophages by plating them at a density of 2-4 x 10^5^ cells/mL in RPMI 1640 with L-glutamine, 10% FBS and 1X pen/strep supplemented with 50 ng/ml M-CSF. Partial media changes were performed every three days after seeding. Macrophages were used for experiments within 7-10 days after seeding.

### Activation of Bone Marrow Derived Macrophages by Lipopolysaccharide

Bone marrow derived macrophages (BMDM) were treated with lipopolysaccharide (LPS) isolated from *Salmonella typhimurium* (Millipore Sigma, Cat # L6143) for one hour at 37° C. For BMDM that were also treated with purified recombinant UMOD, treatment occurred for 30 minutes prior to addition of LPS. Samples were then harvested by scraping and centrifugation prior to lysis and RNA isolation using the RNAqueous-Micro Total RNA Isolation Kit (Invitrogen Life Sciences, Cat #AM1931). Real time PCR was performed as described below.

### Production of Recombinant Uromodulin

A vector to produce recombinant truncated Uromodulin was produced by GenScript. The synthetic sequence of a truncated form of purified human Uromodulin we have used previously^16^ (Supplemental File S1) along with a C-terminal six histidine (6xHis) tag was cloned into the EcoRI/HindIII sites of a GenScript mammalian expression vector. The resulting plasmid was amplified for transcription into the CHO-Express cell line. Recombinant protein was purified from the resulting cell line by GenScript and provided at an endotoxin level of less than 1 EU/mg.

### Western Blot Analysis

Mouse kidneys were lysed in RIPA buffer before Precellys homogenization (Bertin Instruments) according to the manufacturer’s protocol. Samples were separated on a 4-12% Bis-Tris Bolt Gel (Invitrogen Life Technologies) and transferred to a 0.45 µm PVDF membrane. Blots were probed with a goat polyclonal antibody directed against HAVCR1 (KIM-1, R&D Systems, Cat # AF1817) and imaged using Super Signal Femto West Maximum Sensitivity Substrate (ThermoFisher Scientific).

### Real time PCR

RNA extraction from whole kidneys was done using TriReagent (Ambion, Cat. #AM9738) according to manufacturer’s protocol and BMDM cells as described above. Both sample types were reverse transcribed as previously described^14^. Real-time PCR was performed in an Applied Biosystems (AB) ViiA 7 system using TaqMan Gene Expression Assays also from AB. Primers for transcripts of interest are listed in Supplemental File S2. GAPDH was used as a control for all reactions (ThermoFisher Scientific, Cat# 4352339E). Cycling parameters were: 50°C for 2 min, then 95°C for 10 min followed by 40 cycles of 95°C for 15 sec and 60°C for 1 min. Expression values were normalized to endogenous controls and reported as fold change compared to control using the delta-delta CT method, according to the manufacturer’s instructions.

### Statistical Analysis

Values of each experimental group are reported as mean ± standard deviation unless otherwise indicated. All statistical tests were performed using GraphPad Prism software unless otherwise indicated. A two tailed t-test was used to examine the difference in means for continuous data. An F test was used to compare variances; in cases where the variances were significantly different (p < 0.05), a Welch’s correction was applied to the t-test. A paired t-test was used for samples from the same patient collected at different times. The Fisher’s exact test was used to determine differences between categorical variables. Simple linear regressions were used to determine relationships between two continuous variables. Statistical significance was determined at p < 0.05. Statistical outliers were defined using the Tukey’s fences methods. Kaplan-Meier survival curves were analyzed using the Gehan-Breslow-Wilcoxon test to determine statistical significance.

## Results

### Combined spatial protein imaging and single cell transcriptomics to identify early cellular and molecular determinants in ischemia reperfusion injury and increased susceptibility of UMOD^-/-^mice

To understand the impact of UMOD on early molecular and cellular determinants of acute kidney injury (AKI), scRNA-Seq and CODEX (Co-detection by indexing) were performed in a bilateral ischemia reperfusion injury (IRI) model prior to serum creatinine elevation in UMOD^+/+^ and UMOD^-/-^ mice (**Fig. 1b, c**). This timepoint has been used previously in rodent models to identify early predictors of subsequent damage^32,33^. UMOD^-/-^ mice have increased susceptibility to AKI as evidenced by increased serum creatinine versus UMOD^+/+^ 24 hours post injury (peak injury, p = 0.0122, **Fig. 1a**). The 22-minute clamp time induces a mild kidney injury in UMOD^+/+^ mice while the 30-minute clamp time induces severe kidney injury in UMOD^+/+^ mice, similar to what is seen in UMOD^-/-^ mice with a 22-minute clamp time^33,34^.

**Fig. 1.**
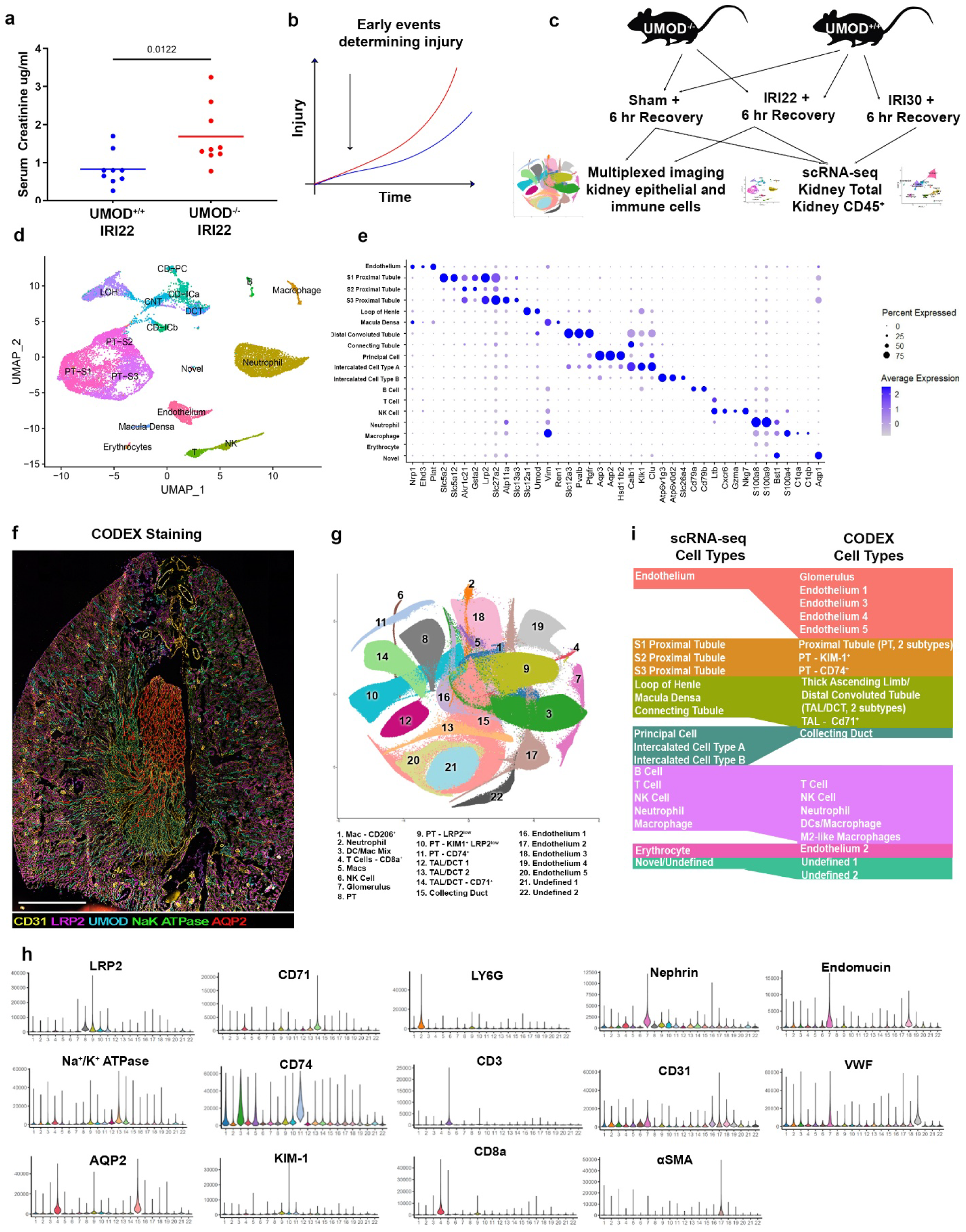
Identification of kidney epithelial, endothelial and immune cells at the single cell level using transcriptomics and spatial protein imaging. **a)** UMOD^-/-^ have increased serum creatinine levels 24 hours following bilateral ischemia reperfusion of the kidneys with 22 minute clamp time (22IRI, n = 9 per group) **b)** Visualization of hypothesis that characterizing earlier timepoints following IRI surgery will identify early molecular events determining later injury **c)** Study design showing contribution of mouse genetic lines (UMOD^-/-^ and UMOD^+/+^) and surgical models to multiplexed imaging and kidney total and Kidney CD45^+^ single cell RNA-sequencing **d)** Clustering of cell types identified in kidney total single cell RNA-sequencing **e)** Marker gene expression in single cell RNA-sequencing clusters **f)** Representative image of whole mouse kidney slice following multiplexed imaging by CODEX **g)** Clustering of cell types identified by CODEX multiplexed imaging **h)** Violin plots of antibody signal from individual cell clusters derived from CODEX multiplexed imaging **i)** Comparison of cell types identified by Kidney Total scRNA-seq and CODEX UMOD – Uromodulin, IRI – ischemia-reperfusion injury, PT-S1 – S1 proximal tubule, PT-S2 – S2 proximal tubule, PT-S3 – S3 proximal tubule, LOH – Loop of Henle, DCT – distal convoluted tubule, CNT – connecting tubule, CD-PC – Collecting Duct – Principal Cell, CD-ICa – Collecting Duct – Intercalated Cell Type A, CD-ICb – Collecting Duct – Intercalated Cell Type B, LRP2 – LDL Receptor Related Protein 2, CD71 – Cluster of Differentiation 71, LY6G – Lymphocyte Antigen 6 Family Member G, CD74 – Cluster of Differentiation 74, CD3 – Cluster of Differentiation 3, CD31 – Cluster of Differentiation 31, VWF – Von Willebrand Factor, AQP2 – Aquaporin 2, KIM-1 – Kidney Injury Molecule 1, CD8a – Cluster of Differentiation 8a, αSMA – α Smooth Muscle Actin

### Single cell spatial protein and transcriptomic landscape of the mouse kidney

A transcriptomic atlas of early injury and UMOD susceptibility was constructed from the single cell transcriptomics data (**Fig. 1d** and **Fig. S1a**). Putative cell types were determined using the top 5 differentially expressed markers (**Fig. 1d, e** and **Fig. S1b**). In parallel, a spatial protein image atlas was built with CODEX imaging using 25 markers (**Table 1 and Fig. S2**) of kidneys from sham surgery and IRI22 mice (**Fig. 1f**). Individual cells were segmented, markers quantitated, and putative cell types of the nephron and immune cells identified (**Figs. 1g and 1h**). Cell-type annotations were validated by back-mapping cell identities onto the images (**Fig. S3**). The scRNA-seq and CODEX identified overlapping cell populations of the nephron and immune system. While scRNA-seq provided deeper cellular phenotyping, CODEX allowed for mapping of cells in their spatial context (**Fig. 1i**).

### Immune cell enriched single cell transcriptomics and spatial proteomics uncover the complexity and spatial zonation of kidney immune cell subtypes

In sham and ischemia surgeries, immune cells were enriched from kidneys by CD45^+^ FACS, subject to scRNA-Seq and cell-types identified based on the top five differentially expressed markers (**Fig S4a-d**)^35^. Transcriptionally defined macrophages and monocytes were further reclustered into five subtypes of macrophages and two distinct populations of monocytes (**Fig. S4d**)^35^. Similarly, immune cell clusters from the CODEX dataset (**Fig. 2a**) were combined and reclustered (**Fig. 2b**) and annotated using immune-specific markers (**Fig. 2c, Fig. S5**). Immune subclusters were mapped back on to the original image to verify marker expression and assignment of cell identities (**Fig. 2d**, identical field from **Fig. 2a**). scRNA-Seq and CODEX identified expected immune cell types and unique subtypes of macrophages (**Fig. S4e**). Based on expression of common markers in the two datasets (CD74, MHCII, CD169 and CD206) scRNA-Seq-based clusters may overlap with CODEX-based clusters (**Fig. S4f**). The spatial context provided by the CODEX dataset highlights a diversity of dendritic cell and macrophage zonation (**Fig. 2e-f, File S3**). Dendritic cells are abundant in the cortex and outer stripe but not in the inner stripe and inner medulla (**Fig. 2f**). In contrast, macrophages have the highest density in the inner stripe of the outer medulla (**Fig. 2i-j**). A macrophage subtype with intermediate level of MHCII expression (Mac (C)) was highly enriched in the inner stripe with low distribution in other kidney regions (**Fig. 2i**). Immunologically quiescent macrophages with low MHCII and CD74 (Mac (E)) were the most abundant subtype and found throughout the kidney and enriched in the inner stripe. A macrophage subtype with intermediate level of MHCII expression (Mac (B)) was found predominantly in the inner stripe. Putative alternatively activated macrophages (Mac 206) were the least abundant and exhibited no zonation (**Fig. 2j, File S3**).

**Fig. 2.**
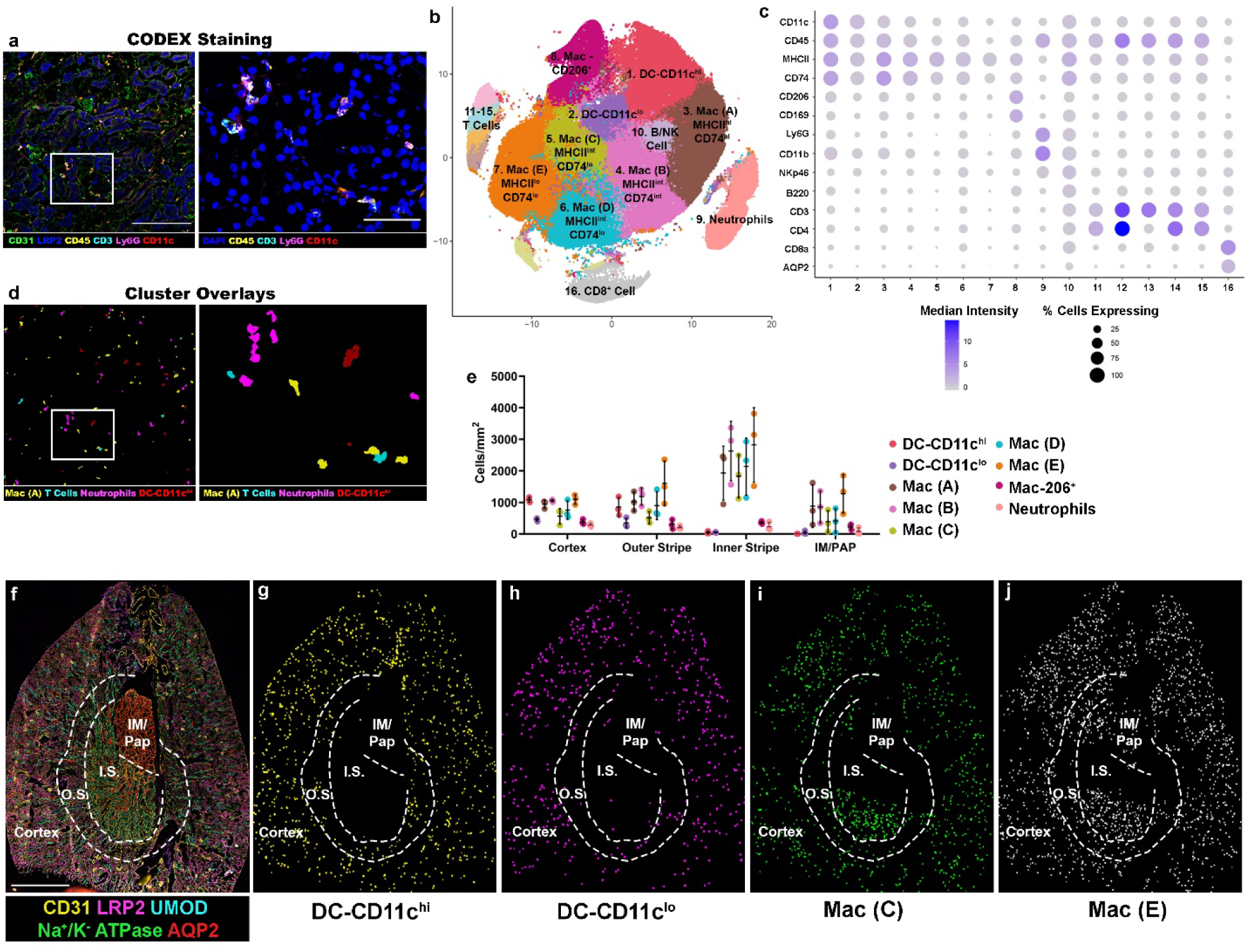
Increased resolution of kidney immune cells at the single cell level using transcriptomics of Cd45^+^-enriched cells and spatial protein imaging. **a)** Representative image of mouse kidney slice following multiplexed imaging by CODEX with selected immune cell markers **b)** Clustering of CD45^+^ cell types identified by CODEX multiplexed imaging **c)** Dotplot of antibody signal from individual CD45^+^ cell clusters derived from CODEX multiplexed imaging **d)** Cluster overlay of CD45^+^ cell clusters onto identical images from (a) **e)** Distribution of myeloid cell type clusters by region in UMOD^+/+^ sham kidney slices (n = 3) **f)** Representative whole kidney slice from UMOD^+/+^ sham kidney with selected markers **g)** Representative image of dendritic cell CD11c^hi^ population in UMOD^+/+^ sham kidney **h)** Representative image of dendritic cell CD11lo^i^ population in UMOD^+/+^ sham kidney **i)** Representative image of macrophage Mac (C) population in UMOD^+/+^ sham kidney **j)** Representative image of macrophage Mac (E) population in UMOD^+/+^ sham kidney UMOD – Uromodulin, DC – dendritic cell, mac - macrophage

### The distal nephron transcriptome responds early to ischemia reperfusion injury

To identify genes that were upregulated with mild (22IRI, 22 minutes ischemia reperfusion injury) and severe (30IRI, 30 minutes IRI) injury in the scRNA-seq analysis of UMOD^+/+^ samples, individual tubule cell types were combined into the proximal tubules, thick ascending limb/distal convoluted tubule and collecting ducts (Principal Cells, Intercalated Cells Types A and B). Differentially expressed transcripts were used to identify unregulated pathways between tubular subsegments and in injury (**File S4**). Transcripts associated with cell adhesion, metabolism and growth^36^, mitochondrial biology, ER/oxidative stress^37,38^, inflammation^39^ and immune signaling^40,41^ (**Fig. 3a**, **File S4**) were upregulated across injury subgroups, suggesting that these are shared tubular epithelium responses to early injury. Individual tubule types also showed specific responses to injury. Specifically, the distal tubular epithelium responds early to injury by up-regulating pathways of wound healing, cell adhesion, and cell migration/cytoskeletal rearrangement. Further, in severe injury (30IRI), apoptosis pathways were up-regulated in the distal nephron concomitant with increases in expression of pro-inflammatory cytokines and specifically cytokine receptor upregulation in the collecting duct (**Fig. 3a-b**, **Fig. 6c-f, File S4**).

**Fig. 3.**
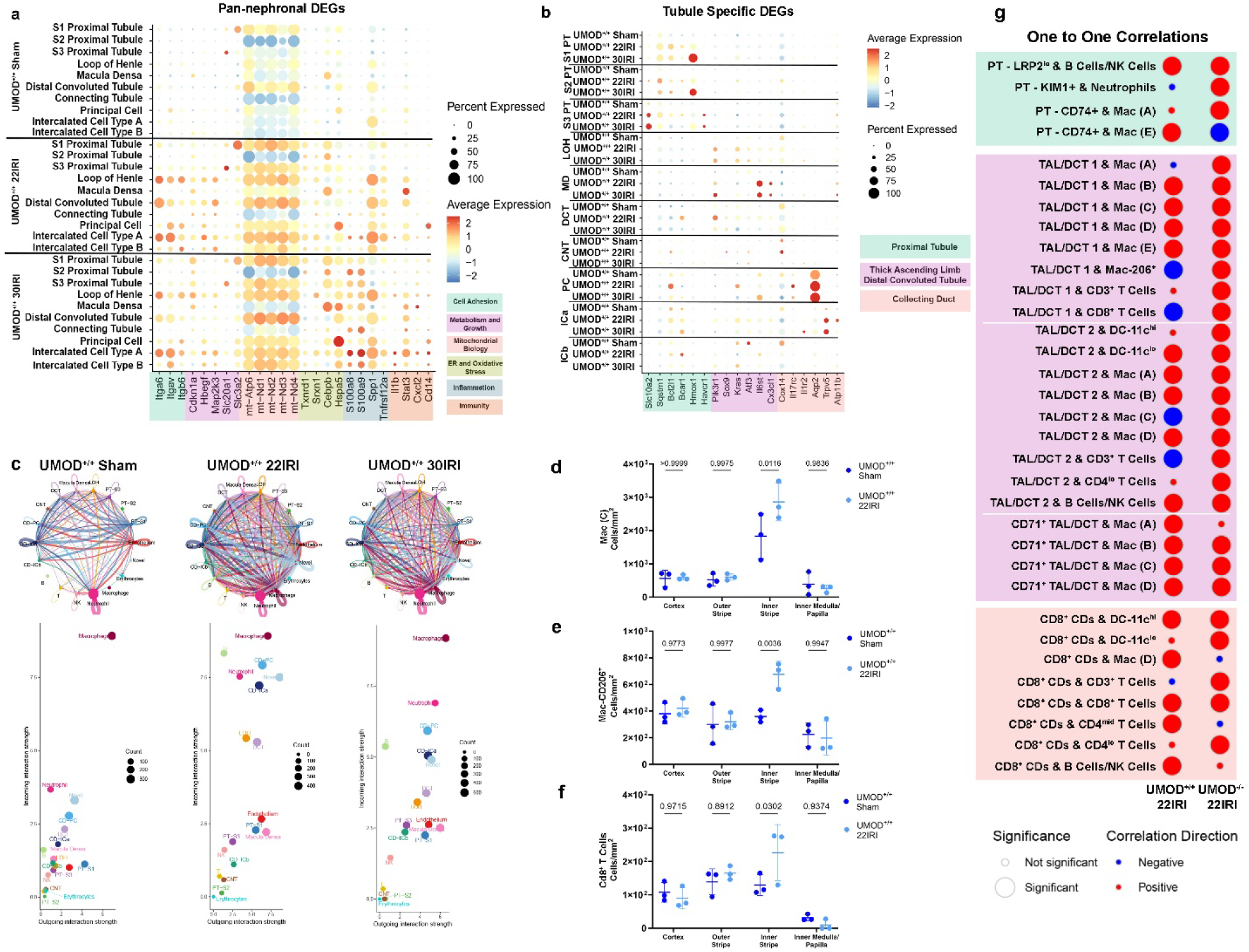
Early ischemic injury to the kidney is marked by shared and unique tubule responses and increased cellular communication between the distal nephron and immune cells. **a)** Increased expression of genes involved in cell adhesion, metabolism and growth, mitochondrial biology, ER and oxidative stress, inflammation and immunity is found with both moderate (22IRI) and severe (30IRI) injury across multiple tubule cell types by scRNA-seq **b)** Increased expression of genes with moderate (22IRI) and severe (30IRI) injury that are unique to proximal tubules, thick ascending limb/distal convoluted tubule and collecting duct by scRNA-seq **c)** (Top) Increased intercellular communication occurs with both moderate (22IRI) and severe (30IRI) injury; (Bottom) Increased intercellular communication in moderate (22IRI) injury is driven by increases in neutrophils, B cells and cells from the distal nephron (CD-PC, CD-ICa, LOH and DCT) which are partially lost with severe injury (30IRI) **d)** Concentration (cells/mm^2^) of Mac (C) population is increased in the inner stripe with moderate injury (22IRI, n = 3 per group) **e)** Concentration (cells/mm^2^) of Mac CD206^+^ population is increased in the inner stripe with moderate injury (22IRI, n = 3 per group) **f)** Concentration (cells/mm^2^) of CD8^+^ T cell population is increased in the inner stripe with moderate injury (22IRI, n = 3 per group) **g)** UMOD^+/+^ 22IRI show shifts in interactions of immune cells with epithelial cells compared to UMOD^+/+^ sham DEG – differentially expressed gene, UMOD – Uromodulin, 22IRI – ischemia-reperfusion injury with 22 minute clamp time, Mac – macrophage, PT – proximal tubule, LRP2 - LDL Receptor Related Protein 2, NK – Natural Killer, KIM1 – Kidney-Injury Molecule 1, CD74 – Cluster of Differentiation 74, TAL – thick ascending limb, DCT – distal convoluted tubule, CD3 – Cluster of Differentiation 3, CD8 – Cluster of Differentiation 8, CD71 – Cluster of Differentiation 71, DC – dendritic cell, CD4 – Cluster of Differentiation 4

### Early cell-cell communication after ischemia reperfusion injury occurs predominantly in the distal nephron

An analysis of immune cell counts in the whole kidney scRNA-seq dataset indicated the proportions of immune cells increase with injury. Specifically, neutrophils increased in numbers and fraction of immune cells (**Fig. S7a,** p < 0.0001). The recruitment of neutrophils in injury was verified *in situ* to the outer and inner stripes of the outer medulla by confocal imaging and CODEX imaging (**Fig. S7b-d**). This suggests immune cells are already actively responding to injury and may be communicating with other kidney cells, consistent with reports from others^42–44^. To test for this possibility, CellChat analysis was performed on the whole kidney transcriptome atlas in UMOD^+/+^ kidneys to predict ligand-receptor interactions. Both with 22IRI and 30IRI there was increased communication as compared to sham (**Fig. 3c**, top panels). While macrophage signaling was the most prominent in all three datasets, 22IRI samples showed increased signaling interactions from B cells and neutrophils. Notably, collecting duct cells (both principal cells and intercalated type A) and the loop of Henle/distal convoluted tubules showed the highest levels of signaling interactions among epithelial cells (**Fig. 3c**, bottom middle panel). Most of the signaling interactions for these cell types showed a partial return to sham conditions in the 30IRI with decreasing incoming interactions, though they still maintained similar outgoing interaction strength as the 22IRI condition. This suggests the increased cell signaling involving the distal nephron, which is partially blunted in 30IRI group, could play a role in the recovery of kidney function seen with 22IRI^34^. Further, specifically in immune cell populations the top activated pathways in immune cells were different in 22IRI versus 30IRI suggesting that the molecular programs activated early on in immune cells could also be important modulators of the severity of injury (**Fig. S8**).

### Ischemia reperfusion leads to early recruitment and association of immune cells to thick ascending limb cells and distal nephron

To quantify the distribution of immune infiltration of the kidney at this early stage of injury a localization analysis was performed (**File S3**). In addition to neutrophil infiltration, there was a significant infiltration of macrophage subtypes (Mac (C) and Mac 206) and CD8^+^ T cells to the inner stripe of the outer medulla, where there is the highest density of TAL cells (**Fig. 3d-f**). Using a cell-centric unbiased neighborhood analysis to quantitatively assess spatial associations between cell types, a total of 96 neighborhood clusters were detected with varying cell type composition and significant spatial associations between cell pairs (**Fig. S9, Fig. 3g**). In UMOD^+/+^ mice, IRI led to significant positive spatial correlation of neutrophils and Mac (A) with injured proximal tubules (PTs). While Mac (A) associated with CD71^+^ TAL/DCT in sham, upon injury Mac (A) shifted to an association with TAL/DCT1. This may be explained by the loss of CD71 expression in TAL cells in the inner stripe with injury (**Fig. S10a-b**). In contrast, Mac (C) and Mac 206 association with TAL/DCT was unique to injury (**Fig. 3g**). New spatial associations of T cell subtypes with TAL/DCT cells, and not PTs, were observed with injury. In addition, a subset of T cells is associated with a novel subpopulation of collecting duct cells expressing the marker CD8a. Cumulatively, these findings suggest that despite increased susceptibility of PTs to kidney injury in the outer stripe of the medulla, early immune-epithelial crosstalk in kidney injury prominently involves the distal nephron and the under-appreciated inner stripe of the medulla.

### Uromodulin deficiency causes early changes to the tubular transcriptome in ischemia reperfusion injury that promote a more severe molecular phenotype

To identify up-regulated genes and molecular pathways with UMOD deficiency (**Fig. 4a-b)** in injury (22IRI for UMOD^-/-^ and UMOD^+/+^**, File S5**) differentially expressed genes in injured UMOD^-/-^ versus injured UMOD^+/+^ mice were identified. Uromodulin deficiency was associated with upregulation of transcripts associated with cell adhesion^45–47^, metabolism and growth^48,49^, mitochondrial biology^50^, stress and inflammation^51–56^, immune signaling^57–59^ and regulated cell death^60–63^ in all tubule types (**Fig. 4a**, **File S5**), suggesting that these are shared responses to UMOD deficiency in early injury. Proximal tubules specifically up-regulated genes involved in metabolism^64–66^ as well as those involved in responding to oxidative stress^67–69^ (**Fig. 4b**, **File S5, Fig. S11a**). The thick ascending limb/distal convoluted tubules upregulated genes involved in cell signaling^70–72^, cell cycling, and the cytoskeleton^73^ (**Fig. 4b**, **File S5, Fig. S11b**). The collecting ducts upregulated genes involved in protein translation^74^ (**Fig. 4b**, **File S5, Fig. S11c**). Given the presence of cell death, stress/inflammation, oxidative stress and signaling genes among those differentially regulated, these findings are consistent with the increased kidney damage seen in UMOD^-/-^ mice with 22IRI (**Fig. S11c**).

**Fig. 4.**
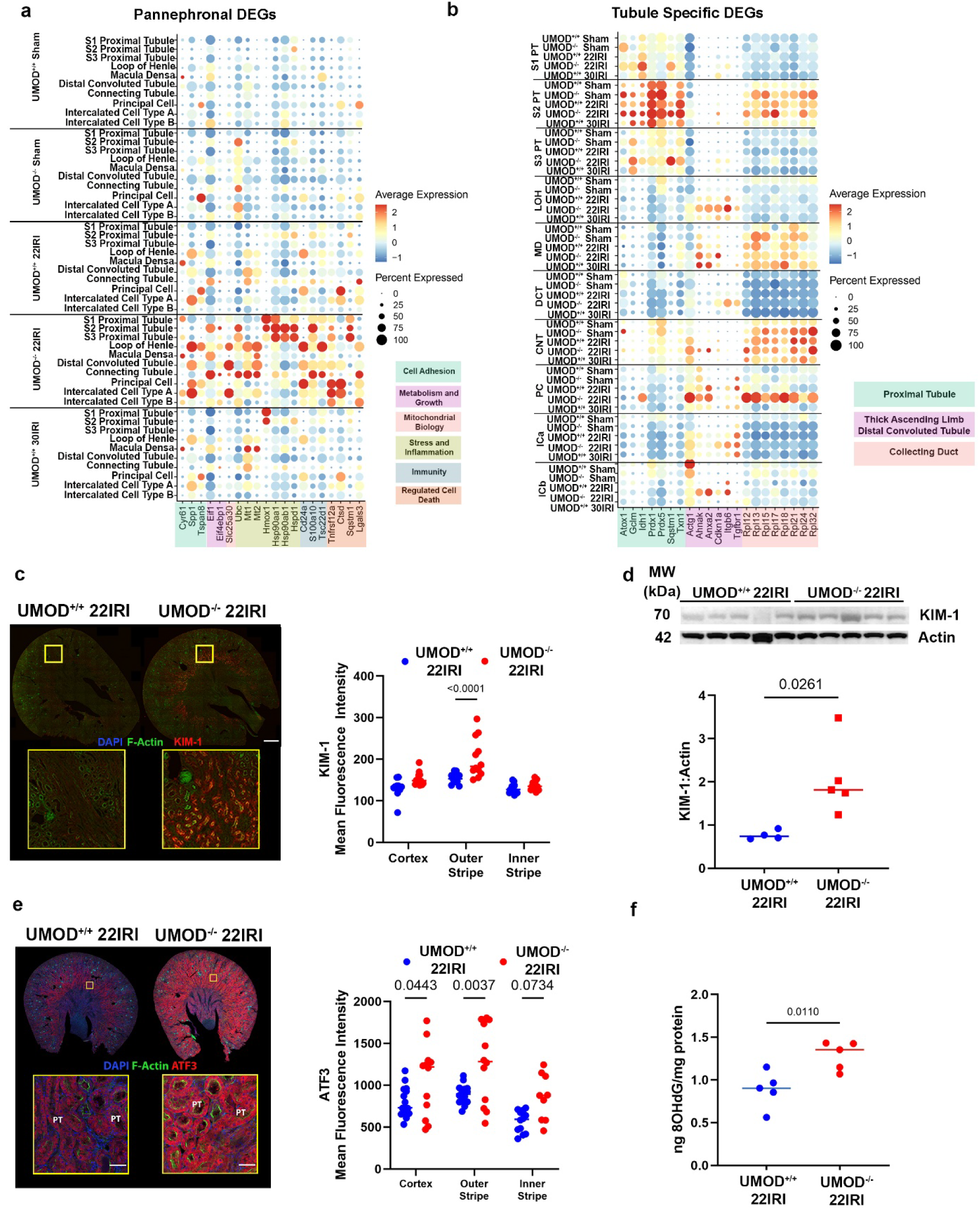
UMOD deficiency leads to early injury marked by upregulation of shared and unique tubule markers, increased expression of injury marker KIM-1 and increased oxidative stress. **a)** Increased expression of genes involved in cell adhesion, metabolism and growth, mitochondrial biology, stress and inflammation, immunity and regulated cell death is seen across multiple tubule types in UMOD^-/-^ 22IRI compared to UMOD^+/+^ 22IRI samples by scRNA-seq **b)** Increased expression of genes in UMOD^-/-^ 22IRI compared to UMOD^+/+^ 22IRI that are unique to proximal tubules, thick ascending limb/distal convoluted tubule and collecting duct by scRNA-seq **c)** Fluorescence intensity of KIM-1 is increased in the outer stripe of UMOD^-/-^ 22IRI compared to UMOD^+/+^ 22IRI mouse kidneys (n = 4 mice per group, 3 fields per mouse per region) **d)** Levels of KIM-1 protein are increased in the kidneys of UMOD^-/-^ 22IRI compared to UMOD^+/+^ 22IRI mice (n = 4-5 per group) **e)** Fluorescence intensity of ATF3 is increased in the cortex and outer stripe of UMOD^-/-^22IRI compared to UMOD^+/+^ 22IRI mouse kidneys (n = 3-4 mice per group, 3 fields per mouse per region) **f)** Levels of oxidative DNA damage (as measured by 8OHdG in tissue lysates normalized to total protein) are increased in UMOD^-/-^ 22IRI compared to UMOD^+/+^ 22IRI mouse kidneys (n = 5 per group) UMOD – Uromodulin, DEG – differentially expressed gene, IRI – ischemia-reperfusion injury, PT-S1 – S1 proximal tubule, PT-S2 – S2 proximal tubule, PT-S3 – S3 proximal tubule, LOH – Loop of Henle, DCT – distal convoluted tubule, CNT – connecting tubule, CD-PC – Collecting Duct – Principal Cell, CD-ICa – Collecting Duct – Intercalated Cell Type A, CD-ICb – Collecting Duct – Intercalated Cell Type B, KIM-1 – Kidney Injury Molecule 1, MW – molecular weight, kDa – kiloDalton, ATF3 – Activating Transcription Factor 3, 8OHdG - 8-hydroxy-2’-deoxyguanosine

To validate and extend these transcriptomic findings, we performed KIM-1 immunofluorescence and imaging of whole mouse kidney sections (**Fig. 4c**) The outer stripe showed increased KIM-1 mean fluorescence intensity in UMOD^-/-^ 22IRI compared to UMOD^+/+^ 22IRI (**Fig. 4c**, p < 0.0001). Immunoblotting for KIM-1 in whole kidney lysates confirmed the increase of KIM-1 expression in injured UMOD^-/-^ kidneys (**Fig. 4d**, p = 0.021). Concomitantly, the expression of the oxidative stress-responsive transcription factor ATF3 was also significantly increased in the kidney cortex and outer stripe at 6 hours in injured kidneys from UMOD^-/-^ compared to UMOD^+/+^ mice (**Fig. 4e**). Consistent with these findings, the levels of 8-Oxo-2’-deoxyguanosine (8-OHdG), a marker for oxidative stress, were also significantly higher in UMOD^-/-^ compared to UMOD^+/+^ mice 6 hours after 22IRI (**Fig. 4f**).

### UMOD deficiency disrupts immune cell recruitment to the inner stripe in early injury

To explore whether UMOD deficiency impacts cellular communication in early injury, we performed CellChat analysis on whole kidney scRNAseq 22IRI data comparing UMOD^+/+^ and UMOD^-/-^. While the magnitude of signaling from the distal nephron is increased in UMOD^-/-^ mice, the signaling from macrophages decreased in UMOD^-/-^ kidneys post injury (**Fig. 5a**). In contrast, a Reactome pathway analysis on the combined mononuclear phagocytes (macrophages, dendritic cells, monocytes) from CD45^+^ scRNA seq demonstrated an increased activation of genes involved in immune activation, such as the type II interferon response, lymphocyte activation, antigen presentation and cell-cell interactions in UMOD^-/-^ 22IRI versus UMOD^+/+^ 22IRI mice (**Fig. 5b**). This increased activation was also seen in other immune populations (**Fig. S12**). Together, these results suggest that Uromodulin may play a pivotal role in regulating immune system activation in response to injury through an interaction with mononuclear phagocytes.

**Fig. 5.**
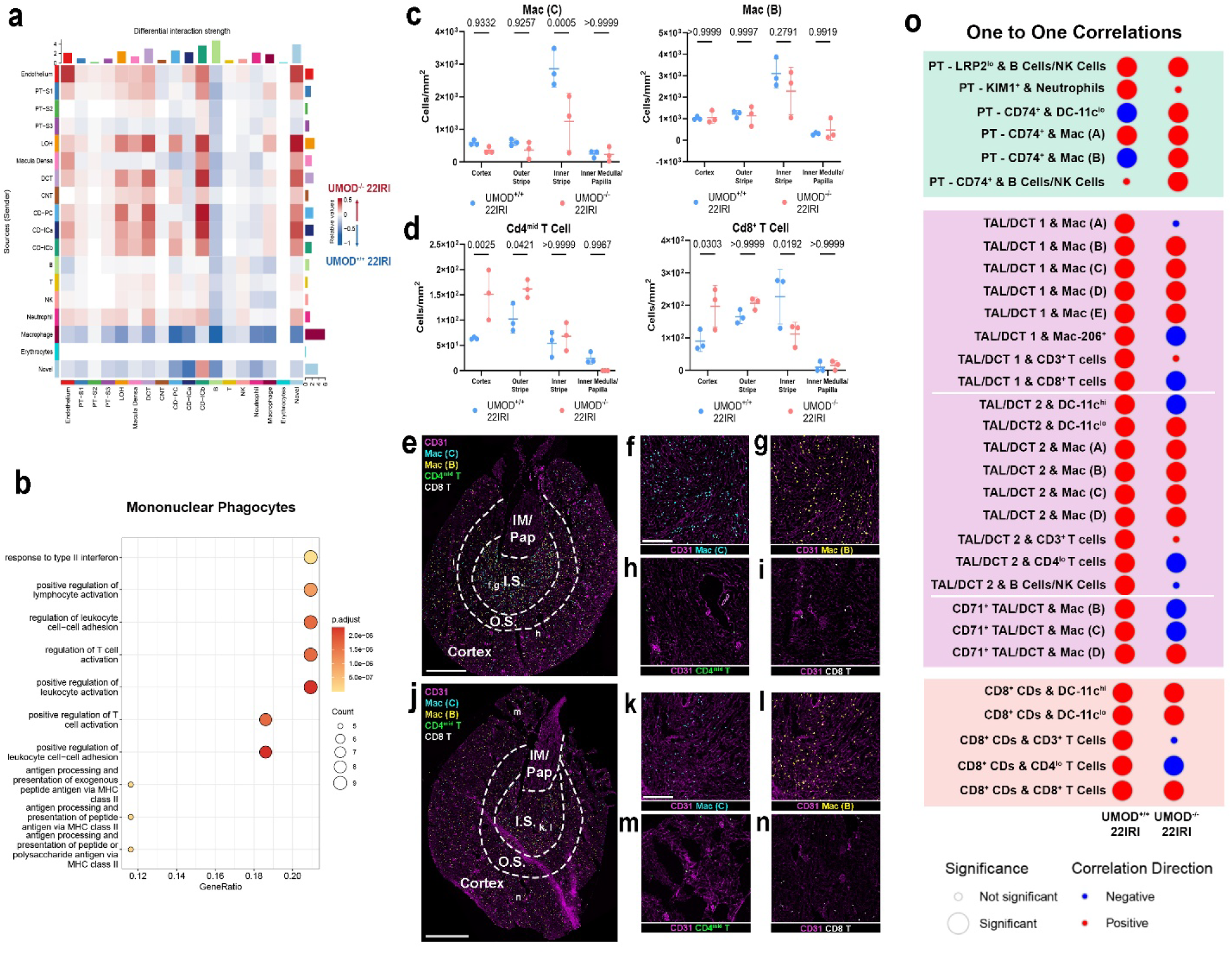
UMOD deficiency disrupts macrophage and T cell zonation in the kidney and increases proinflammatory signaling pathways. **a)** CellChat analysis of scRNA-seq shows an increase in signaling from the proximal and distal nephron but decreased signaling from macrophages and a novel cell type with Uromodulin deficiency in samples with moderate (22IRI) injury **b)** Mononuclear phagocytes (macrophages, dendritic cells, monocytes) from UMOD^-/-^ 22IRI have increased signaling in pathways related to type II interferon, lymphocyte/T cell activation, leukocyte cell-cell adhesion and antigen presentation compared to those from UMOD^+/+^ 22IRI **c)** (Left) Concentration (cells/mm^2^) of Mac (C) population is decreased in the inner stripe of UMOD^-/-^ mice compared to UMOD^+/+^ mice with moderate injury (22IRI, n = 3 per group) (Right) Concentration (cells/mm^2^) of Mac (B) population is trending toward a decrease in the inner stripe of UMOD^-/-^ mice compared to UMOD^+/+^ mice with moderate injury (22IRI, n = 3 per group) **d)** (Left) Concentration (cells/mm^2^) of CD4^mid^ T cell population is increased in the cortex and outer stripe of UMOD^-/-^ mice compared to UMOD^+/+^ mice with moderate injury (22IRI, n = 3 per group) (Right) Concentration (cells/mm^2^) of CD8^+^ T cell population is increased in the cortex and decreased in the inner stripe of UMOD^-/-^ mice compared to UMOD^+/+^ mice with moderate injury (22IRI, n = 3 per group) **e-n)** Representative distribution of Mac (C), Mac (B), CD4mid T cells and CD8+ T Cells in UMOD^+/+^ 22IRI (top) and UMOD^-/-^ 22IRI (bottom) kidneys **o)** UMOD^-/-^ 22IRI show increased recruitment of immune cell types to the proximal tubule and decreased recruitment to the TAL/DCT and collecting ducts compared to UMOD^+/+^ 22IRI UMOD – Uromodulin, DEG – differentially expressed gene, IRI – ischemia-reperfusion injury, PT-S1 – S1 proximal tubule, PT-S2 – S2 proximal tubule, PT-S3 – S3 proximal tubule, LOH – Loop of Henle, DCT – distal convoluted tubule, CNT – connecting tubule, CD-PC – Collecting Duct – Principal Cell, CD-ICa – Collecting Duct – Intercalated Cell Type A, CD-ICb – Collecting Duct – Intercalated Cell Type B, Mac – Macrophage, CD4 – Cluster of Differentiation 4, CD8 – Cluster of Differentiation 8, IM – inner medulla, Pap – papilla, LRP2 - LDL Receptor Related Protein 2, NK – Natural Killer, KIM1 – Kidney-Injury Molecule 1, CD74 – Cluster of Differentiation 74, TAL – thick ascending limb, DCT – distal convoluted tubule, CD3 – Cluster of Differentiation 3, CD71 – Cluster of Differentiation 71, DC – dendritic cell

To determine how these changes might impact immune cell recruitment and immune-epithelial interactions, we performed a regional immune cell analysis to quantify the effect of Uromodulin deficiency on immune infiltration of the kidney. Among myeloid cells, there was a significant reduction of Mac (C) density in the outer stripe of UMOD^-/-^ 22IRI mice. Mac (B) had a similar trend but was not significant (**Fig. 5c**). There was also an increase in the cell density of CD4^+^ T cells in the cortex and outer stripe areas of the kidney (Cd4^mid^ population shown, but the increase was also seen in Cd4^low^ T Cells, **Fig. 5d, left**). A shift of CD8^+^ T cell abundance towards the cortex was also observed in UMOD^-/-^ kidneys (**Fig. 5d, right**). Selected images validating this regional analysis are shown (**Fig. 5e-n**). We next compared specific neighborhood pairs to study the impact of Uromodulin deficiency on immune-epithelial spatial interactions during injury (**Fig. 5o**). We found that new associations now occurred between immune cells (DC-11c^hi^, Mac (B) and B/NK Cell) and proximal tubules in UMOD^-/-^ mice. Additionally, spatial associations between immune cells and the distal nephron (particularly TAL/DCT) seen in UMOD^+/+^ kidneys were not present in UMOD^-/-^ kidneys (Mac (A), DC-11c^hi^, Mac (B), Mac (C), M2 and various T cell subtypes). This change cannot be solely explained by the decreased macrophage homing to the inner stripe (which primarily affects Mac (C)).

Taken together, these findings suggest that with injury prone to repair, Uromodulin promotes homing and confinement of immune cells to the inner stripe. This could protect areas susceptible to injury, such as the outer medulla (could extend to the medullary rays in the cortex), from uncontrolled activation of the immune system. However, in UMOD deficiency, this immune zonation and confinement is altered and there is a shift in immune cell distribution towards more diffuse involvement of the outer stripe and cortex.

### UMOD inhibits Nlrc4 expression in macrophages which drives the alternative activation of the inflammasome

Since *UMOD^-/-^* kidneys are more susceptible to injury, we investigated the expression of genes associated with cell death using qPCR in injured kidneys from *UMOD^-/-^* and *UMOD^+/+^* mice. We found no differences between the two genotypes for genes associated with apoptosis (*Bak1*, *Bax*, **Fig. 6a**) or necroptosis (*Mlkl*, *Ripk1*, **Fig. 6b**). However, there was increased expression of the inflammasome-associated IL-1β at both the transcript and protein levels (**Fig. 6c**). To determine the potential cell source of excess Il-1β in UMOD^-/-^ kidneys, we explored the scRNA-seq data. *Il1b* is predominantly expressed by neutrophils in all groups and in macrophages with injury and/or Uromodulin deficiency (**Fig. S13, Fig. 6d**). Consistently, *Nlrp3*, a pattern recognition receptor (PRR) for the inflammasome IL-1β, is also expressed primarily in neutrophils and, to a lesser extent, in the macrophages. In contrast to *Il1b*, *Nlrp3* appeared to be reduced in UMOD^-/-^ neutrophils, making it unlikely that NLRP3 inflammasomes alone are responsible for the increased Il-1β in this setting (**Fig. S13, Fig. 6d**). Consistent with this, a second PRR, *Nlrc4*, was predominantly expressed in the macrophages of UMOD^-/-^ mice in both sham and IRI conditions, suggesting that NLRC4 inflammasomes in macrophages are a second source of IL-1β in the UMOD^-/-^ mice (**Fig. S13, Fig. 6d**). The scRNA-seq findings of elevated *Nlrc4* but not *Nlrp3* expression in UMOD^-/-^ 22IRI mice compared to UMOD^+/+^ 22IRI were confirmed by qPCR of whole kidney RNA (**Fig. 6h**). We also observed increased IL-23 expression in UMOD^-/-^ mice, consistent with increased inflammasome activation leading to expression of IL-23 (**Fig. 6i**).

**Fig. 6.**
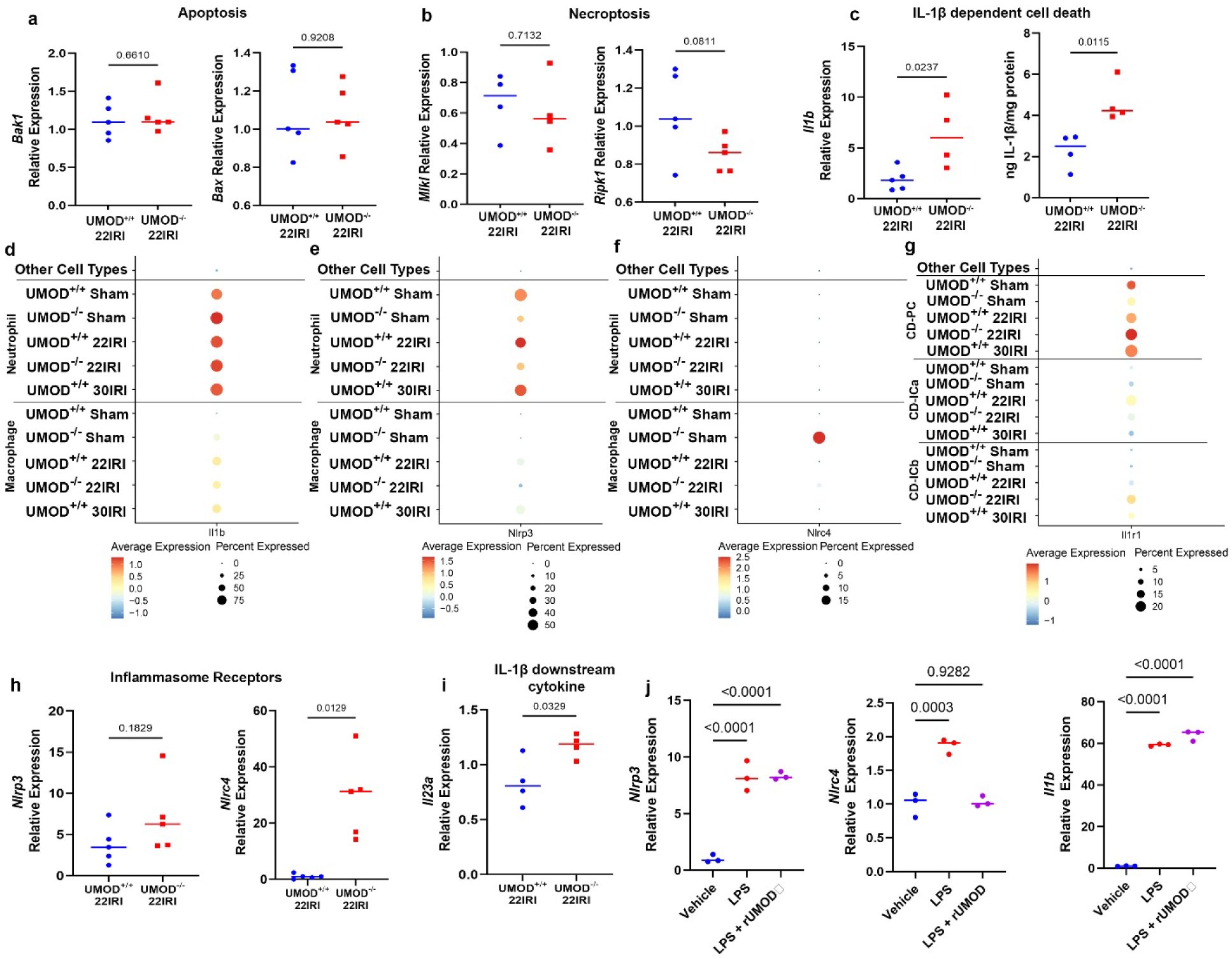
UMOD deficiency increases IL-1β-dependent cell death via expression of the non-canonical inflammasome *Nlrc4*. **a)** There is no difference in expression of apoptosis markers *Bak1* and *Bax* in UMOD^-/-^22IRI compared to UMOD^+/+^ 22IRI mouse kidneys (n = 5 per group) **b)** There is no difference in expression of necroptosis markers *Mlkl* and *Ripk1* in UMOD^-/-^22IRI compared to UMOD^+/+^ 22IRI mouse kidneys (n = 5 per group) **c)** Expression of the *Il1b* transcript as well as levels of IL-1β protein normalized to total protein are increased in UMOD^-/-^ 22IRI compared to UMOD^+/+^ 22IRI mouse kidneys (n = 5 per group) **d)** *Il1b* is primarily expressed by neutrophils and macrophages in all genotype/surgery groups by scRNA-sequencing. **e)** The canonical inflammasome sensor *Nlrp3* is expressed in both UMOD^+/+^ and UMOD^-/-^groups **f)** The non-canonical sensor *Nlrc4* is exclusively expressed in UMOD^-/-^ groups. **g)** A receptor for IL-1β is expressed in distal nephron segments (*Il1r1*). **h)** There is no difference in expression of canonical inflammasome sensor *Nlrp3* in UMOD^-/-^22IRI compared to UMOD^+/+^ 22IRI mouse kidneys (n = 5 per group) but expression of the non-canonical inflammasome sensor *Nlrc4* transcript is increased in UMOD^-/-^ 22IRI compared to UMOD^+/+^ 22IRI mouse kidneys (n = 5 per group) **i)** Expression of the IL-1β-dependent *Il23a* transcript is increased in UMOD^-/-^ 22IRI compared to UMOD^+/+^ 22IRI mouse kidneys (n = 5 per group) **j)** Pre-treatment with 100 ng/ml of recombinant truncated Uromodulin (rUMOD) does not prevent induction of *Nlrp3* expression by *Salmonella typhimurium* lipopolysaccharide (LPS) **k)** Pre-treatment with 100 ng/mL of recombinant truncated Uromodulin (rUMOD) prevents induction of *Nlrc4* expression by *Salmonella typhimurium* lipopolysaccharide (LPS) **l)** Pre-treatment with 100 ng/mL of recombinant truncated Uromodulin (rUMOD) does not prevent induction of *Il1b* expression by *Salmonella typhimurium* lipopolysaccharide (LPS) UMOD – Uromodulin, IRI – ischemia-reperfusion injury, PT-S1 – S1 proximal tubule, PT-S2 – S2 proximal tubule, PT-S3 – S3 proximal tubule, LOH – Loop of Henle, DCT – distal convoluted tubule, CNT – connecting tubule, CD-PC – Collecting Duct – Principal Cell, CD-ICa – Collecting Duct – Intercalated Cell Type A, CD-ICb – Collecting Duct – Intercalated Cell Type B

To verify these findings *in vitro*, we differentiated macrophages from bone marrow cells isolated from UMOD^+/+^ mice and treated with *S. typhimurium* lipopolysaccharide (LPS) to induce expression of *Nlrp3* and *Nlrc4*. Pre-treatment with 100 ng/mL recombinant truncated UMOD (rUMOD) did not impact *Nlrp3* induction by LPS (**Fig. 6j, left**) but blocked *Nlrc4* induction (**Fig. 6j, middle**). Expression of *Il1b* increased significantly with LPS treatment irrespective of rUMOD pre-treatment (**Fig. 6j, right**).

To determine potential targets of IL-1β, we studied the expression of the IL-1 receptors. *Il1r1* was primarily expressed in principal cells of the collecting ducts and in macula densa cells with injury (**Fig. S13, Fig. 6g**). A lower level of expression was observed in endothelial cells and intercalated type A and B cells of the collecting ducts (**Fig. S13, 6g**). *Il1r2* was primarily expressed in neutrophils, with a low level of expression in macrophages with injury (**Fig. S13**). Therefore, the target of excess IL-1β could be other myeloid cells and/or specific epithelial cells such as collecting duct cells. Similarly, when examining the expression of *Il23a* in the scRNA seq data, it was expressed in neutrophils (in UMOD^-/-^ mice or UMOD^+/+^ injury groups), and in intercalated type A cells of the collecting ducts with UMOD^-/-^ injury (**Fig. S13**). Taken together, these findings suggest that the excess inflammation (for example increased IL-23 production) in UMOD deficiency during injury could be downstream of IL-1β pro-inflammatory activation of immune cells and collecting duct cells.

### UMOD deficiency leads to increased expression of Cd8a by Collecting Duct Cells expressing Il1r1

To better understand the impact of IL-1β signaling on collecting duct cells in the injured UMOD^-/-^kidneys we examined transcriptomic and proteomic expression profiles in collecting duct epithelial cells. We detected a cluster of CD8-expressing collecting duct (CD) cells in the CODEX data (CD8^+^, AQP2^+^ and CD3^-^, **Fig. S14**). These CD8^+^ CD cells were contiguous with normal collecting duct cells within the cortex (**Fig. 7a-c, Fig. S15**). While these cells were observed in all the groups included, they were significantly more abundant in UMOD^-/-^ mice (**Fig. 7d**). To assess whether this may be due to endogenous expression rather than protein uptake, we examined the whole kidney scRNA-seq dataset. Collecting duct cell types (principal cells and intercalated cell type A) also express *Cd8a* at the transcriptional level (**Fig. 7e**). Within collecting duct cells, the highest expression of *Cd8a* was observed in 1) uninjured UMOD^-/-^principal vells (PC) and intercalated cell type A (ICa) cells, and 2) in injured UMOD^-/-^ ICa (**Fig. 7f**). When PC and ICa were subclustered, we identified: two sub-clusters of ICa, CD-ICa_3 and CD-ICa_4 (**Fig. 7g**),and one sub-cluster of PC, CD-PC_0 (**Fig. 7h**), expressing *Cd8a*. Compared to *Cd8a*-negative clusters, these clusters also showed elevated expression of *Il1r1*, the receptor for Il-1β, consistent with activation of the inflammasome (**Fig. 7g-h**). These cells uniquely expressed *Gata2*, a transcription factor that triggers an inflammatory phenotype of collecting duct cells (**Fig. 7g-h**). Interestingly, *Il23a* was expressed in only a minority of CD-ICa cells that were separate from the *Cd8a*^+^ CD cells (**Fig. S16**), which suggests that the contribution of inflammasome-mediated IL-23 expression in UMOD^-/-^ mice is mostly driven by immune cells.

**Fig. 7.**
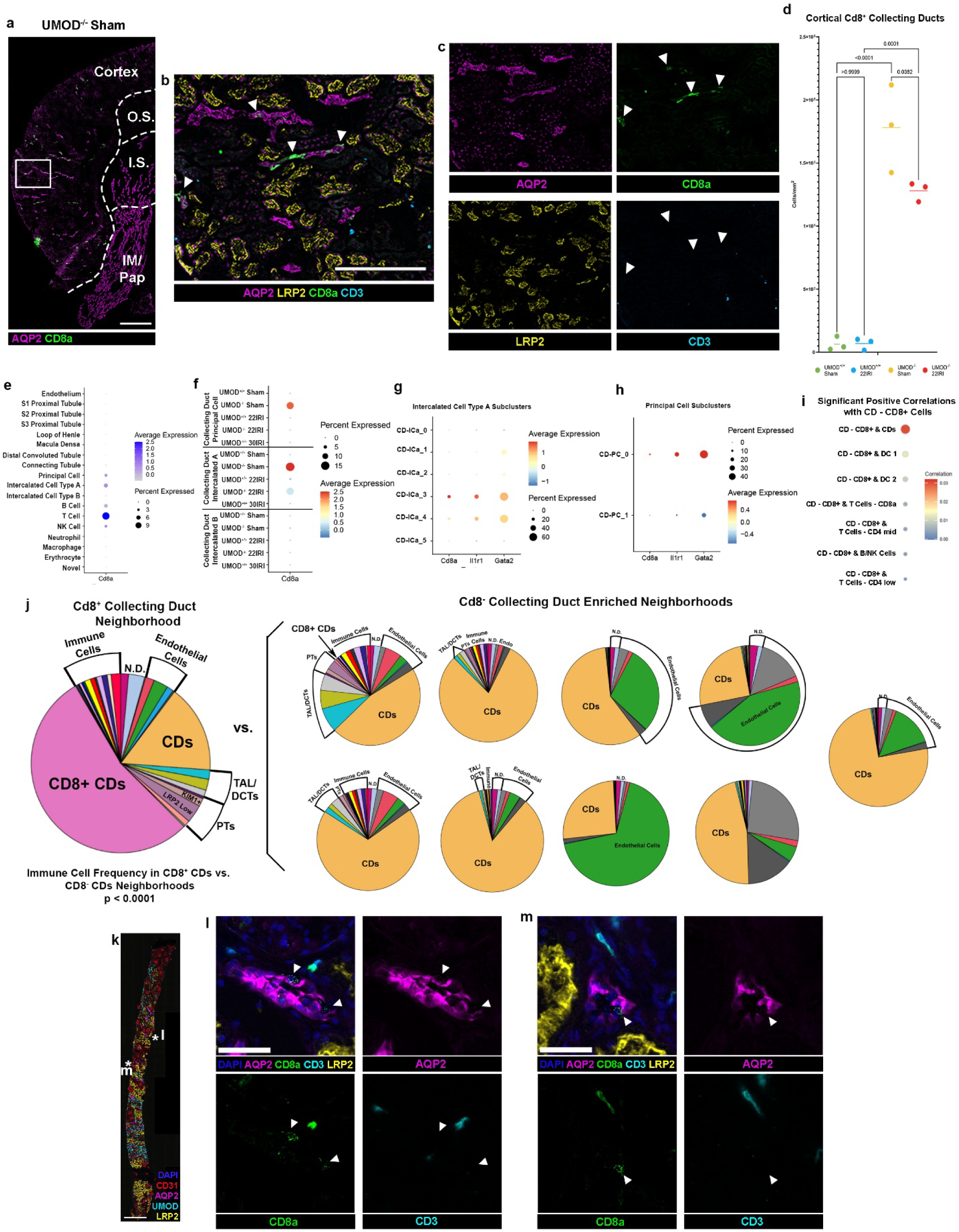
UMOD deficiency increases expression of T Cell Marker *Cd8a* by Collecting Duct cells, which is associated with immune cell localization. **a)** Representative image of CD8a^+^, AQP2^+^ cells in UMOD^-/-^ sham kidney **b)** Magnification of (a) with additional markers for markers for proximal tubules (LRP2) and T cells (CD3) **c)** Individual panels for (B) **d)** Concentration (cells/mm^2^) of CD8^+^ collecting ducts in the cortex increases with UMOD deficiency and decreases in UMOD knockout mice with injury **e)** *CD8a* is expressed in Principal and intercalated Type A cells of the collecting duct in addition to immune cells **f)** *CD8a* expression is highest in UMOD^-/-^ sham Principal and Intercalated Type A cells, but is also expressed in Intercalated Type A cells in UMOD^-/-^ 22IRI kidneys **g)** Two subclusters of Intercalated Type A cells express *CD8a* and these subclusters also express the IL-1β receptor *Ilr1* and the pro-inflammatory transcription factor *Gata2* **h)** One subcluster of Principal Cells expresses *Cd8a* and this subclusters also expresses the IL-1β receptor *Ilr1* and the pro-inflammatory transcription factor *Gata2* **i)** CD8^+^ Collecting Ducts are positively correlated with normal collecting ducts as well as multiple immune cell subtypes **j)** CD8^+^ Collecting Duct Neighborhoods (left) have a higher immune cell frequency than neighborhoods primarily composed of Cd8^-^ collecting ducts **k)** Identification of CD8+ CD3-cells in proximity to AQP2+ collecting ducts in human biopsies from patients with acute kidney injury (AKI) **l)** Magnification of selected area from (k) **m)** Magnification of selected area from (k) UMOD – Uromodulin, AQP2 – aquaporin 2, LRP2 - LDL Receptor Related Protein 2, CD8a – Cluster of Differentiation 8a, CD3 – Cluster of Differentation 3, IRI – Ischemia-reperfusion injury, CD-PC – Collecting Duct – Principal Cell, CD-ICa – Collecting Duct – Intercalated Cell Type A, CD-ICb – Collecting Duct – Intercalated Cell Type B, CD – Collecting Duct, DC – Dendritic Cell, N.D. - not determined, TAL – thick ascending limb, DCT – distal convoluted tubule, Endo – endothelial cells, KIM1 – Kidney Injuyr Molecule –, PT – proximal tubule, CD31 – Cluster of Differentation 31

To extend and validate the pro-inflammatory signature of Cd8^+^ CD cells, we used pairwise neighborhood analysis in the CODEX data to show that these cells were spatially associated with several immune cells in the tissue (**Fig. 7i**). Furthermore, the composition of neighborhood niches (**Fig. S9**) showed that the CD8^+^ collecting duct neighborhood niche also included CD8^-^collecting duct cells immune cells, endothelial cells, TAL/DCT and proximal tubule cell clusters (**Fig. 7j**). The immune cell frequency in the CD8+ collecting duct neighborhood was significantly higher than in CD8^-^ collecting duct neighborhoods (p < 0.0001, **Fig. 7j**), further highlighting the specificity of the association of these subtypes of collecting ducts to immune cells.

Finally, the presence of CD8^+^ collecting duct cells was validated in the CODEX datasets in patients with kidney disease from the Kidney Precision Medicine Projects (**Fig. 7k-m**), thereby extended the potential clinical relevance of our findings to humans.

## Discussion

In this study, we combined single cell transcriptomics and multiplexed large scale spatial protein imaging to define the molecular events that occur early on, *in situ*, in the evolution of kidney injury. The spatial organization of cellular and molecular effectors likely governs the course of AKI and its severity. We also investigated the role of UMOD in this setting, which is known to increase the severity of AKI in the kidney through its role in modulating immune-epithelial crosstalk.

Within 6 hours after ischemia, kidney tubular cells throughout the entire nephron respond to injury by communicating with each other and with immune cells, predominantly neutrophils and macrophages. Many of the observed signaling pathways are common, but each segment also displays a unique molecular response. Notably, the distal nephron including the TAL segments and collecting ducts appear the most active in cell-cell communication. Such communication was altered with longer ischemia clamp time associated with severe injury, suggesting that an appropriate response to injury that is associated with recovery involves a significant output from the distal nephron. Interestingly, this epithelial-immune molecular dialogue was also associated with zonation of specific infiltrating macrophages and Cd8^+^ T cells to the inner stripe of the outer medulla, which is a site away from the more susceptible outer stripe. This could suggest that an adaptive response to injury involves early guiding of some of the infiltrating macrophages and T cells to the inner stripe. We propose that this process of spatial immune diversion is needed to control the immune response and steer it away from vulnerable sites of injury.

UMOD deficiency swayed the molecular phenotype of individual cell segments towards more injury which manifested by increased injury markers such as KIM-1 in proximal tubules and more oxidative stress. This is consistent with our previous findings and can anchor the current results with previous work^14,17^. UMOD deficiency altered the magnitude and the quality of cell-cell communication, particularly in macrophages. Despite a reduction in macrophage infiltration to the inner stripe, which is consistent with our previous findings^16^, the signaling from macrophages output was reduced and was polarized towards a pro-inflammatory phenotype. Furthermore, there was a loss of association of many macrophage subtypes and T cells with TAL cells, and shifting towards the cortex and outer stripe, supporting a role for UMOD in immune zonation and spatial diversion.

On a molecular level, UMOD^-/-^ mice have aberrant increased expression of the alternative inflammasome associated molecule *Nlrc4*, which is consistent with increased IL-1β levels seen in the kidneys of these mice. The transcriptomic findings from the kidneys were also validated *ex vivo* and supported a direct role for UMOD in inhibiting *Nlrc4* expression on macrophages. Therefore, not only can UMOD facilitate the recruitment of macrophages to the inner stripe, but it also tempers the immune system after injury by inhibiting the inflammasome activation through downregulating *Nlrc4* expression in macrophages, which leads to reduced release of IL-1β. Augmented release of IL-1β can have impactful downstream effects on targets cells expressing the IL-1 receptor, such as neutrophils and macrophages which express interleukin 1 receptor type 2 (*Il1r2*). These cells exhibit heightened inflammatory signaling, as shown by their transcriptomic profile from kidneys of UMOD^-/-^ mice. In addition, neutrophils have the highest expression of *Il23a*, which is significantly elevated during injury in UMOD^-/-^ kidneys. Therefore, enhanced expression of *Il23a* could be a downstream effect of inflammasome activation. Although we have previously shown using regional transcriptomics that S3 segments could have induction of *Il23a* expression with UMOD deficiency^15^, those studies were not as comprehensive as single cell RNA sequencing and did not include cells such as neutrophils or collecting duct cells. Furthermore, collecting duct cells also have high expression of interleukin 1 receptor type 1 (*Il1r1*), suggesting that they could also be a target of IL-1β.

Indeed, UMOD deficiency caused a significant increase in the abundance of collecting duct cells expressing the T cell marker *Cd8a.* The expression of CD8 was demonstrated at the protein and RNA levels, thereby validating the endogenous expression of CD8 by this subpopulation and not an uptake of this protein. The absence of CD3 expression also excludes the possibility that the signal is emanating from T cells infiltrating the collecting ducts. The demonstration of these cells in human kidney biopsies infers an important function of this cell subtype in human kidney disease. These *Cd8a*^+^ collecting duct cells expressed *Il1r1*, suggesting they are likely responsive to the increased Il-1β, consistent with their increased prevalence in UMOD^-/-^ mice. *Cd8a*^+^ CD cells were detected as subtypes of principal and intercalated cells. The recent recognition of transitional cells expressing both IC and PC markers particularly in pathophysiological states raises the possibility that CD8+ CDs may be part of this transitional state, and that inflammasome signaling could play a role in this pleiotropy.

Although *Cd8a*^+^ CDs do not express high levels of *Il23a*, this cell subtype has high expression of *Gata2*, a transcription factor implicated in inducing a pro-inflammatory phenotype in collecting duct cells. Yu *et al* showed that inhibiting Gata2 in collecting duct cells ameliorated AKI by reducing inflammatory cytokine production^75^. A recent study by Takai and colleagues using expression and epigenetic chromatin profiling demonstrated that Gata2 binds and induces the expression of key inflammatory loci such as C-X-C motif ligand 10 (*Cxcl10*), vascular cell adhesion molecule-1 (*Vcam1*) and colony-stimulating factor 1 (*Csf1*)^76^. Furthermore, Gata2 has a functional interaction with activator protein (AP-1)^77^, a key transcription factor that regulates inflammation and stress kinase response^78^. The spatial protein imaging data further highlights the pro-inflammatory phenotype of these cells, as compared to other collecting duct cell types, by demonstrating localized immune cell infiltration and enrichment of immune niches around these cells.

The studies presented here put forward important findings that are relevant to the understanding of the kidney response and adaptation during early injury and the role of Uromodulin. Previous studies have outlined the importance of infiltrating immune cells such as T cells and neutrophils as early responders (reviewed here^79^), however there are no previous comprehensive data on the spatial distribution. The demonstration of immune zonation for macrophages and T cells and the role of Uromodulin in enhancing this role is an important finding of this study, especially in the context of immune diversion. Interestingly, neutrophils partially escape this zonation, which is consistent with their role in the pathogenesis in early development of injury^80^. Although we did not have spatial data on IRI 30 minutes, data from UMOD^-/-^ kidneys suggest that escaping this zonation, which was facilitated by UMOD deficiency, is linked to increased severity of injury. This study highlights the importance of investigating the spatial changes in additional studies that link severity and time course of adaptive vs. maladaptive repair to changes in immune cell distribution within the kidney.

An important contribution of this work is showing the global response to injury, particularly the extent of cell communication from the distal nephron, which could be key in orchestrating the response and course of kidney injury. The communication of the distal nephron appears to be a key effector in epithelial – immune crosstalk. Although proximal tubules, and especially S3 segments, are more sensitive to injury, our data suggest that the entire nephron, especially TAL and distal segments, displays an organized response. We propose that a breakdown of this organized response occurs with severe injury or in high-risk states such as Uromodulin deficiency. This in turn will initiate loss of protective mechanisms leading to injury at the most vulnerable sites, and a partial collapse of regulated immune zonation that is meant to temper and organize the immune response.

Other important findings highlighted by this work is the existence of various levels of epithelial-immune communications and the importance of Uromodulin in regulating this crosstalk through inhibiting alternative activation of the inflammasome signaling in macrophages. Although the role of inflammasome activation and IL-1β is becoming more established in kidney injury, the role of NLRC4-driven activation in kidney injury is less well recognized. Our findings suggest that activation of this pathway could contribute significantly to polarization of the macrophage response towards a pro-inflammatory phenotype and that immunomodulators, such as Uromodulin, could use this pathway to fine tune and regulate this injury-associated immune response.

On a technical level, our study demonstrates how spatial protein imaging can be used effectively, in coordination with transcriptomics data, to unravel molecular events *in situ* that govern pathology in an organ. Previous work by other groups have used spatial transcriptomics to define molecular events at the cell level during kidney injury^81–83^. Our current study extends these efforts to spatial proteomics and showcases how outputs could be integrated with single cell transcriptomics to yield important biological insights. Similar efforts using human kidney tissue specimens in the kidney^23^ and other organs^84^ are being used to link spatial molecular events to unravel the trajectory of disease.

Our study has limitations that are worth discussing. Due to the focus on an ischemic model of injury, this study may be limited in its generalizability to nephrotoxic type of AKI. However, many of our important findings such as distribution landscape of the immune system in non-injured kidneys, the impact of UMOD deficiency on immune recruitment and macrophage *Nlrc4* expression and the characterization of CD8^+^ CD cells are independent of the type of injury and will likely have broader applications. Furthermore, this study was designed *a priori* to focus on a single timepoint to study the spatial and molecular changes in early injury. Incorporating similar studies at later timepoints will be needed to define trajectory of changes. However, this will require larger scale of collaborative efforts and resources (workforce, computational, financial etc.). We opted to relay our findings on this important early timepoint and establish an analytical pipeline that could be incorporated in such complimentary future efforts. Some of our findings require additional investigation beyond the scope of this work, including unraveling the molecular mechanism by which circulating Uromodulin inhibits *Nlrc4* expression in macrophages, further expanding on the function of CD8-expressing collecting ducts in the kidney and exploring the hypothesis that UMOD acts as a chemoattractant to attract immune cells broadly to the kidney and specifically to the thick ascending limb.

In conclusion, using integrated transcriptomic and multiplexed spatial protein imaging, we uncovered cellular and molecular changes *in situ* that underlie early stages of kidney injury and defined the role of Uromodulin in this setting. Our findings support a role for this protein in spatially confining the immune system around TAL cells in the inner stripe away from the vulnerable outer stripe. Uromodulin also inhibits *Nlrc4* expression on macrophages, and IL-1β-driven macrophage-epithelial crosstalk that could induce collecting duct cells towards more inflammatory signaling.

## Data Availability Statement

All sequencing data were analyzed using the GRCm38 (mm10) reference genome. For data from the KPMP atlas, results are based on data generated by the Kidney Precision Medicine Project, accessed on Dec 15, 2025 at https://www.kpmp.org. These raw imaging data are under controlled access (human data) as they are potentially identifiable and can be accessed from the following respective sources: 1) KPMP data has been deposited at https://atlas.kpmp.org/repository/ and can be requested and made available by signing a data use agreement (DUA) with KPMP. KPMP will respond to initial data requests within 12–36 h and provide data up to one month after DUA has been signed. Manuscripts resulting from KPMP controlled access data are requested to go through the KPMP Publications and Presentations Committee which reviews and approves manuscripts every two weeks. Any analysis resulting from KPMP data may be published or shared so long as it does not re-identify KPMP participants. Source data are provided with this paper. For the imaging data: the source data for all the main and supplementary figures (confocal imaging, CODEX, and scRNA-seq) will be made available in one or more Zenodo repositories upon acceptance of this manuscript.

## Author Contributions

ARS, SW, TEA, KL designed, conducted, analyzed and interpreted experiments and wrote the manuscript. AN and RM designed, analyzed and interpreted experiments and edited the manuscript. DW, CJG and SK designed and conducted experiments.

## Funding

This work was supported by The National Institute of Diabetes Digestive and Kidney Disease (NIDDK: 1R01DK111651, U54DK137328 to TME), R00 award (R00DK127216) to KL, Veterans Affairs Merit Awards (I01BX003935 and I21BX006519 to TME), an award from the Dialysis Clinic Inc. to TME, and Takeda Science Foundation for AN. The Kidney Precision Medicine Project (KPMP) is supported by the National Institute of Diabetes and Digestive and Kidney Diseases (NIDDK) through the following grants: U01DK133081, U01DK133091, U01DK133092, U01DK133093, U01DK133095, U01DK133097, U01DK114866, U01DK114908, U01DK133090, U01DK133113, U01DK133766, U01DK133768, U01DK114907, U01DK114920, U01DK114923, U01DK114933, U24DK114886, UH3DK114926, UH3DK114861, UH3DK114915, and UH3DK114937. We gratefully acknowledge the essential contributions of our patient participants and the support of the American public through their tax dollars. The content is solely the responsibility of the authors and does not necessarily represent the official views of the National Institutes of Health.

## Acknowledgment

The authors acknowledge the University of Michigan Medical School Central Biorepository (RRID:SCR_026845) for providing biospecimen storage, management, and distribution services in support of the research reported in this publication/grant application/presentation. We acknowledge the Indiana University Medical Genomics Core for performing RNA sequencing, the Indiana University O’Brien Center of Advanced Microscopy (U54DK137328) and the University of Alabama O’Brien Core for serum creatinine measurements. The authors acknowledge Dr. Pierre Dagher for helpful discussions and review of the manuscript.

## Supplemental Data

Raw and processed sequencing data will be made available through the Gene Expression Omnibus repository when the manuscript is accepted for publication.

## Notes

### Competing Interest Statement

The authors have declared no competing interest.

