## Supplemental Figures for "Uromodulin promotes immune zonation and inhibits alternative inflammasome-mediated activation of immune-to-collecting duct inflammatory signaling in early acute kidney injury"

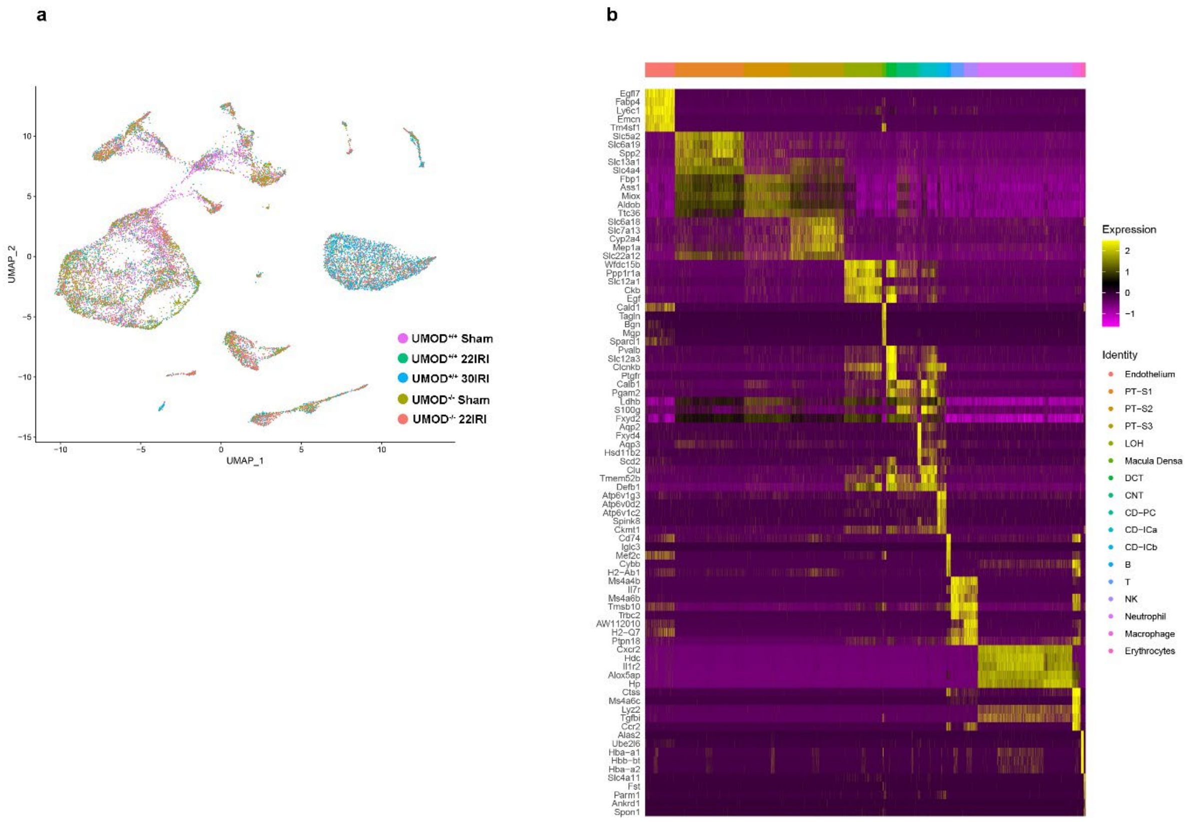


**Fig.S1.** Cell clustering for whole kidney scRNA-seq

**a)** Dimensionality reduction plot of cells from Figure 1d (whole kidney scRNA-seq experiment) split by experimental group

**b)** Heatmap of top 5 marker genes for each cell cluster shown in (a)

UMOD – Uromodulin, IRI – ischemia-reperfusion injury, PT-S1 – S1 proximal tubule, PT-S2 – S2 proximal tubule, PT-S3 – S3 proximal tubule, LOH – Loop of Henle, DCT – distal convoluted tubule, CNT – connecting tubule, CD-PC – Collecting Duct – Principal Cell, CD-ICa – Collecting Duct – Intercalated Cell Type A, CD-ICb – Collecting Duct – Intercalated Cell Type B

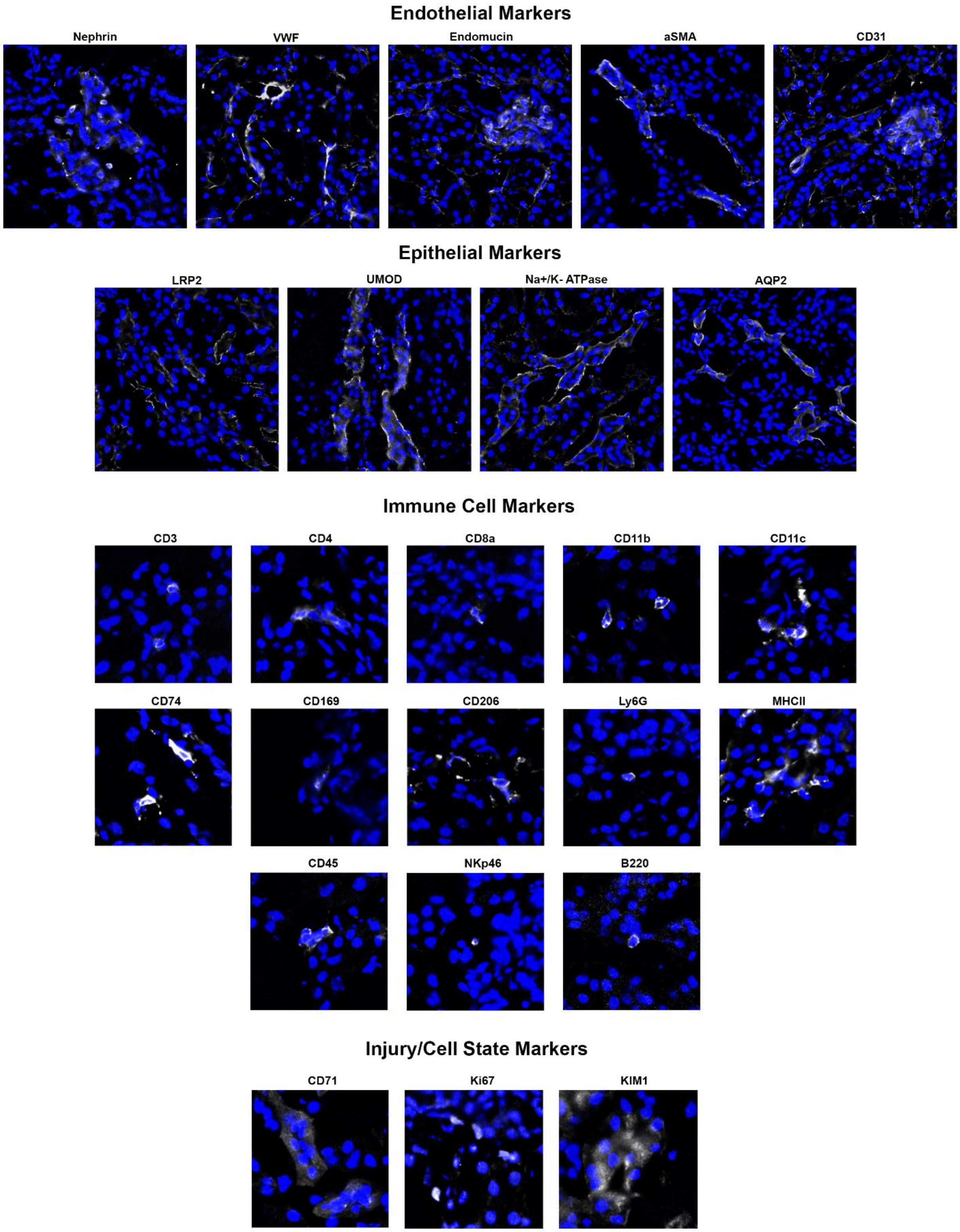


**Fig.S2.** Representative images for each antibody used in the CODEX panel

VWF – Von Willebrand Factor, αSMA – α Smooth Muscle Actin, CD – Cluster of Differentiation, LRP2 – LDL Receptor Related Protein 2, UMOD – Uromodulin, AQP2 – Aquaporin 2, LY6G – Lymphocyte Antigen 6 Family Member G, MHCII – Major Histocompatibility Complex Class II, NKp46 – NK cell p46-related protein, B220 – B-cell member of the T200 glycoprotein family, Ki67 – Antigen Kiel 67, KIM-1 – Kidney Injury Molecule 1

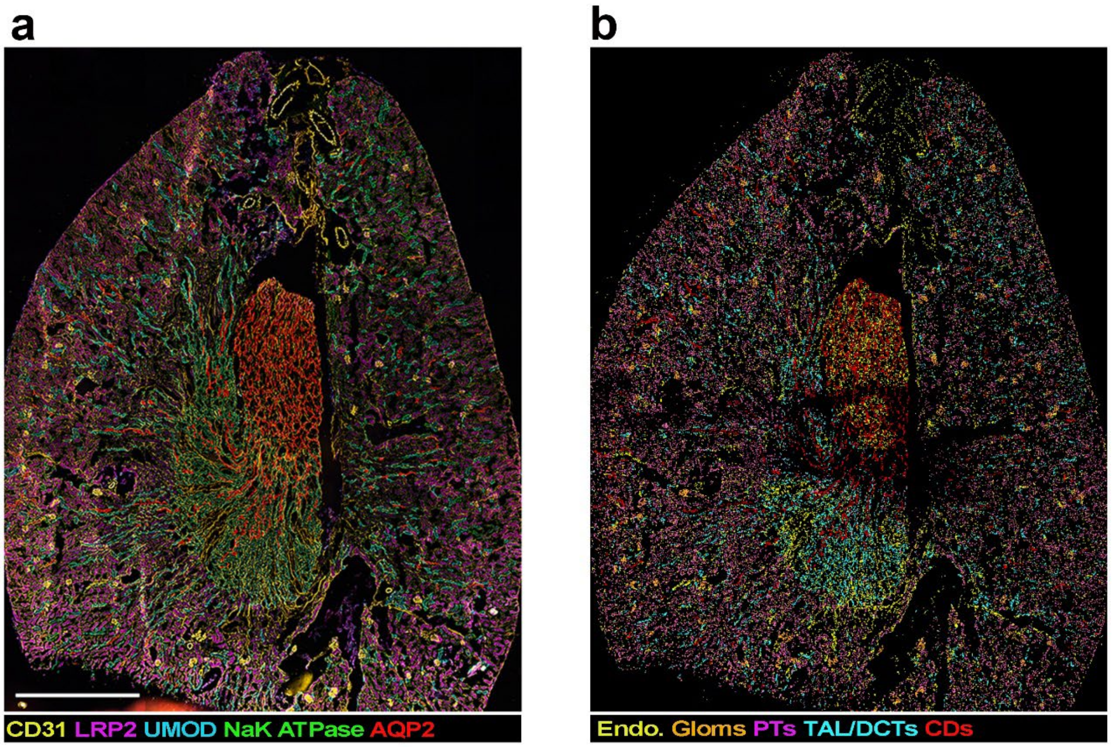


**Fig.S3.** Mapping of CODEX clusters onto representative kidney image

- **a)** CODEX image of a representative kidney slice with selected markers used to identity endothelial and tubular cell types
- **b)** Mapping of CODEX clusters back onto the same kidney slice from (a) shows that endothelial and tubular cell clusters localize to expected structures within the kidney

CD31 – Cluster of Differentiation 31, LRP2 – LDL Receptor Related Protein 2, UMOD – Uromodulin, AQP2 – Aquaporin 2, Endo – endothelial cell clusters, Gloms – glomerular cell clusters, PTs - proximal tubule cell clusters, TAL/ DCTs – thick ascending limb of the loop of Henle and distal convoluted tubule cell clusters, CDs – Collecting Duct cell clusters

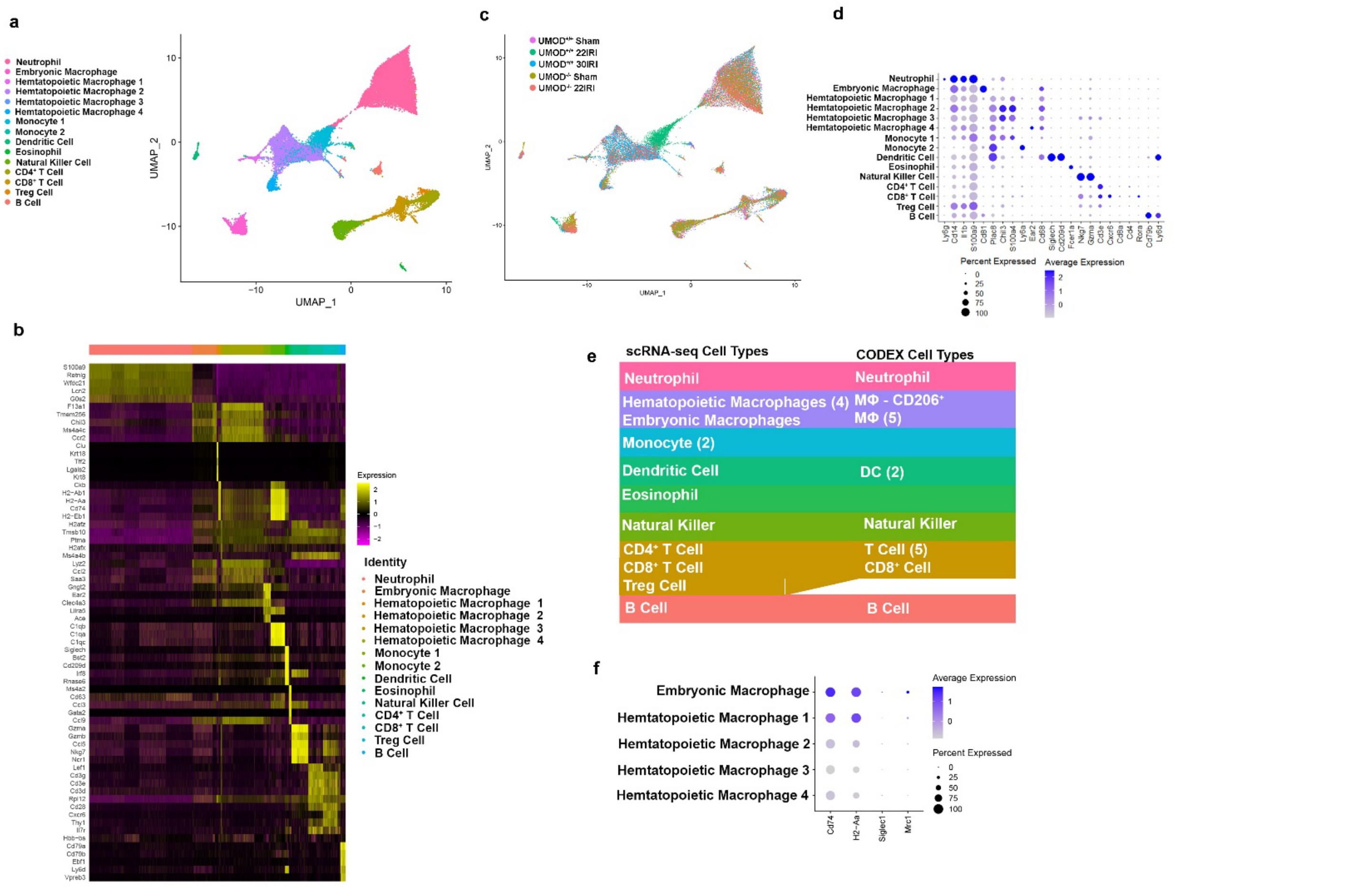

**Fig. S4.** Identification of kidney immune cells at the single cell level using enriched single cell transcriptomics

- **a)** Clustering of cell types identified in kidney Cd45$^+$ immune cell enriched single cell RNA-sequencing
- **b)** Heatmap of top 5 marker genes for each cell cluster shown in (a)
- **c)** Dimensionality reduction plot of cells from (a) split by experimental group
- **d)** Marker gene expression in single cell RNA-sequencing clusters
- **e)** Comparison of cell types identified by Kidney Cd45$^+$ immune cell enriched scRNA-seq and Cd45$^+$ cells in CODEX
- **f)** Marker gene expression in macrophage groups

UMOD – Uromodulin, IRI – ischemia-reperfusion injury, CD – Cluster of Differentiation, LY6G – Lymphocyte Antigen 6 Family Member G, Treg cell – Regulatory T cell

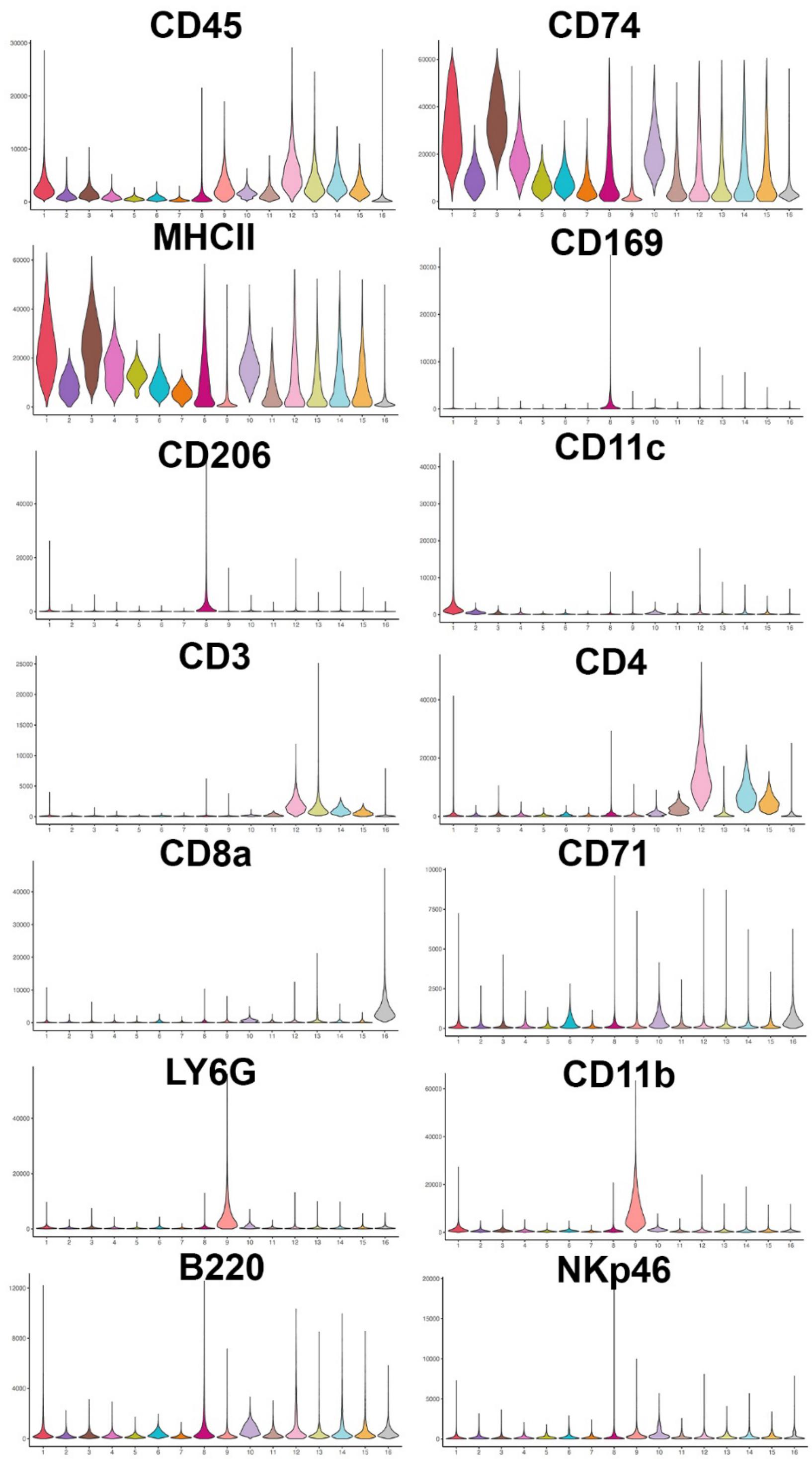


**Fig.S5.** Violin plots of relative expression for each antibody marker in CODEX clusters

CD– Cluster of Differentiation, LY6G – Lymphocyte Antigen 6 Family Member G, MHCII – Major Histocompatibility Complex Class II, NKp46 – NK cell p46-related protein, B220 – B-cell member of the T200 glycoprotein family, Ki67 – Antigen Kiel 67, KIM-1 – Kidney Injury Molecule 1

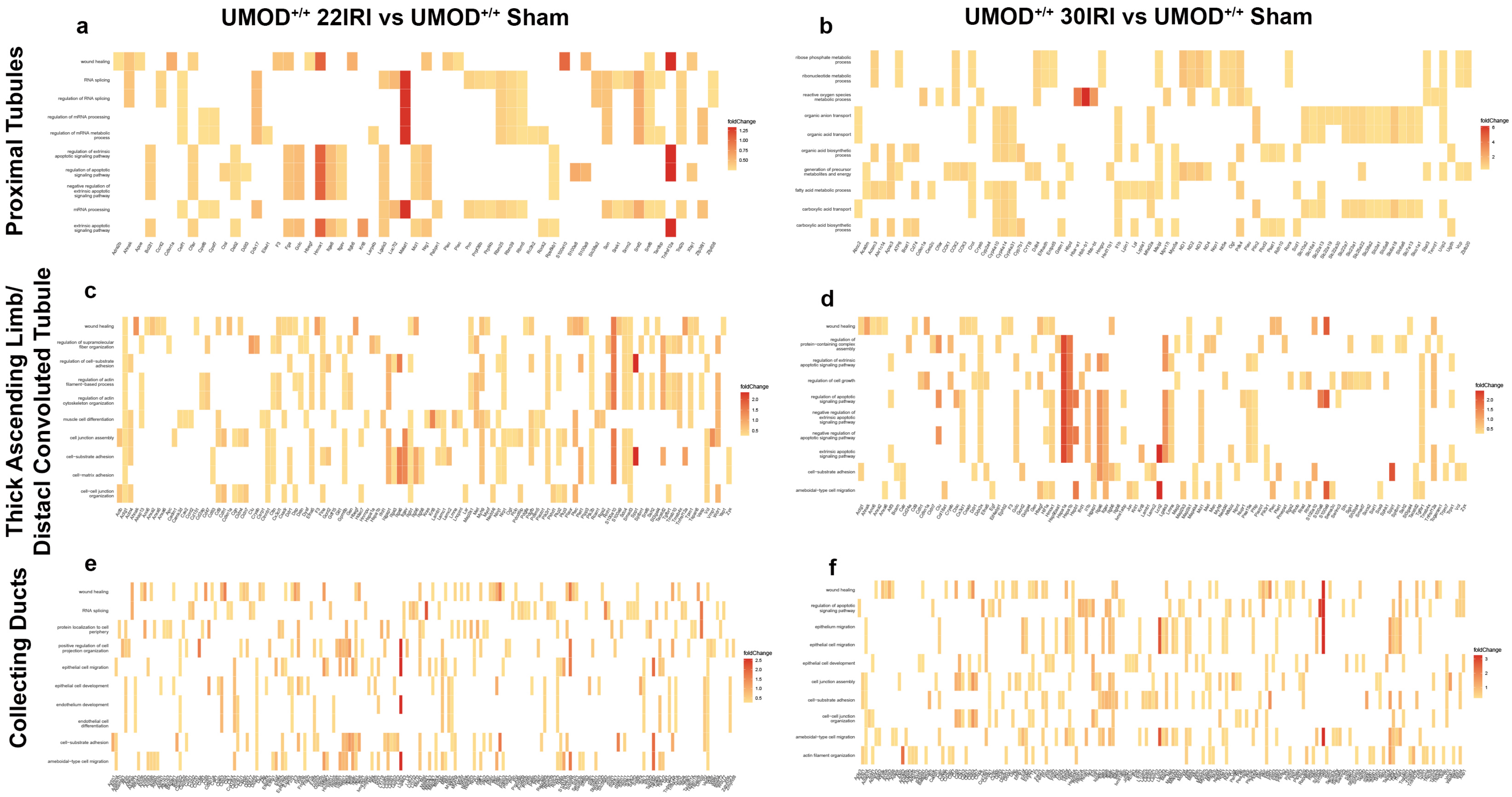


**Fig. S6.** Identification of injury regulated genes and pathways in nephron subsegments

**a-b)** Differentially regulated genes contributing to the identification of the top 10 differentially regulated pathways in proximal tubules with moderate (22IRI, a) and severe (30IRI, b) injury compared to sham

**c-d)** Differentially regulated genes contributing to the identification of the top 10 differentially regulated pathways in thick ascending limb/distal convoluted tubules with moderate (22IRI, c) and severe (30IRI, d) injury compared to sham

**e-f)** Differentially regulated genes contributing to the identification of the top 10 differentially regulated pathways in collecting ducts with moderate (22IRI, a) and severe (30IRI, b) injury compared to sham

UMOD – Uromodulin, IRI – ischemia-reperfusion injury

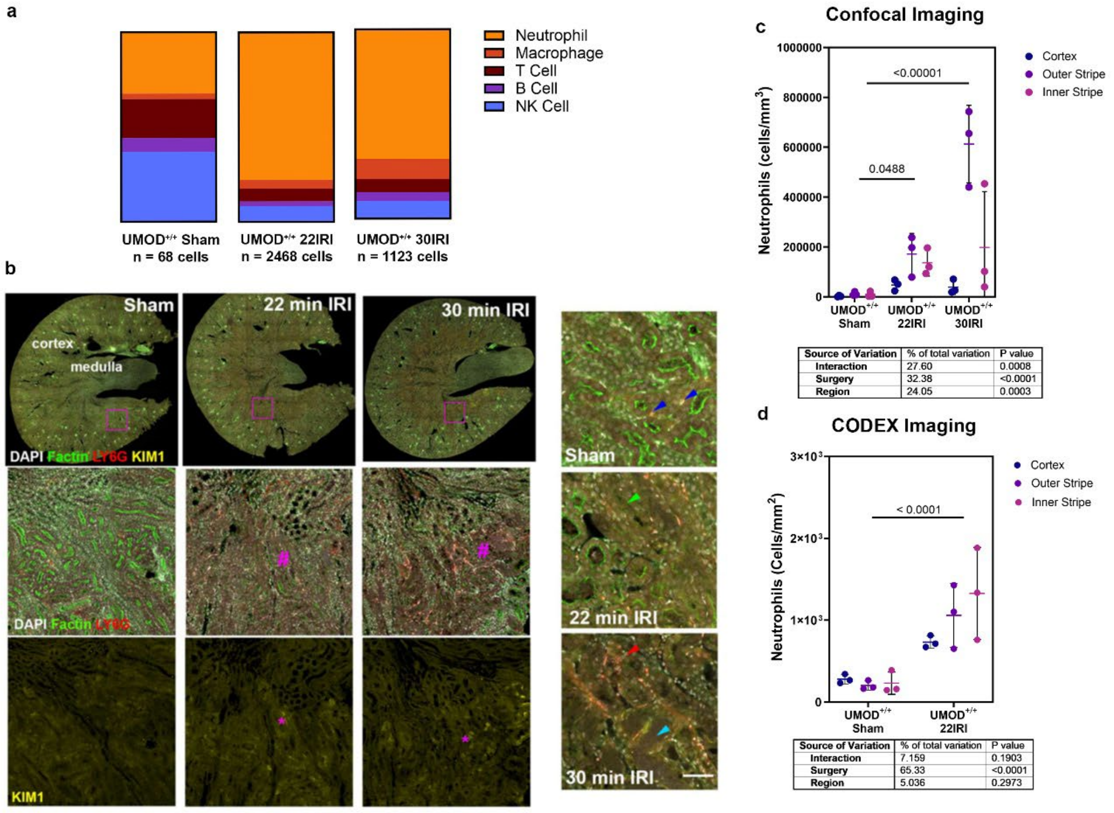


**Fig. S7.** Neutrophils are recruited to the kidney early following ischemia-reperfusion injury

**a)** Distribution of selected immune cells within whole kidney scRNA-seq datasets for UMOD$^{+/+}$ groups showed increased number and proportion of neutrophils

**b)** Representative confocal imaging of LY6G$^{+}$ neutrophils in whole kidney slices from UMOD$^{+/+}$ animals 6 hours after undergoing sham, 22IRI and 30IRI surgeries shows increased recruitment of neutrophils as injury increases which is particularly prominent in the outer stripe of severely injured (30IRI) animals

**c)** Regional distribution of the neutrophil cluster in CODEX data shows increased numbers of neutrophils of injured animals

UMOD – Uromodulin, IRI – ischemia-reperfusion injury, CD – Cluster of Differentiation, LY6G – Lymphocyte Antigen 6 Family Member G, Treg cell – Regulatory T cel

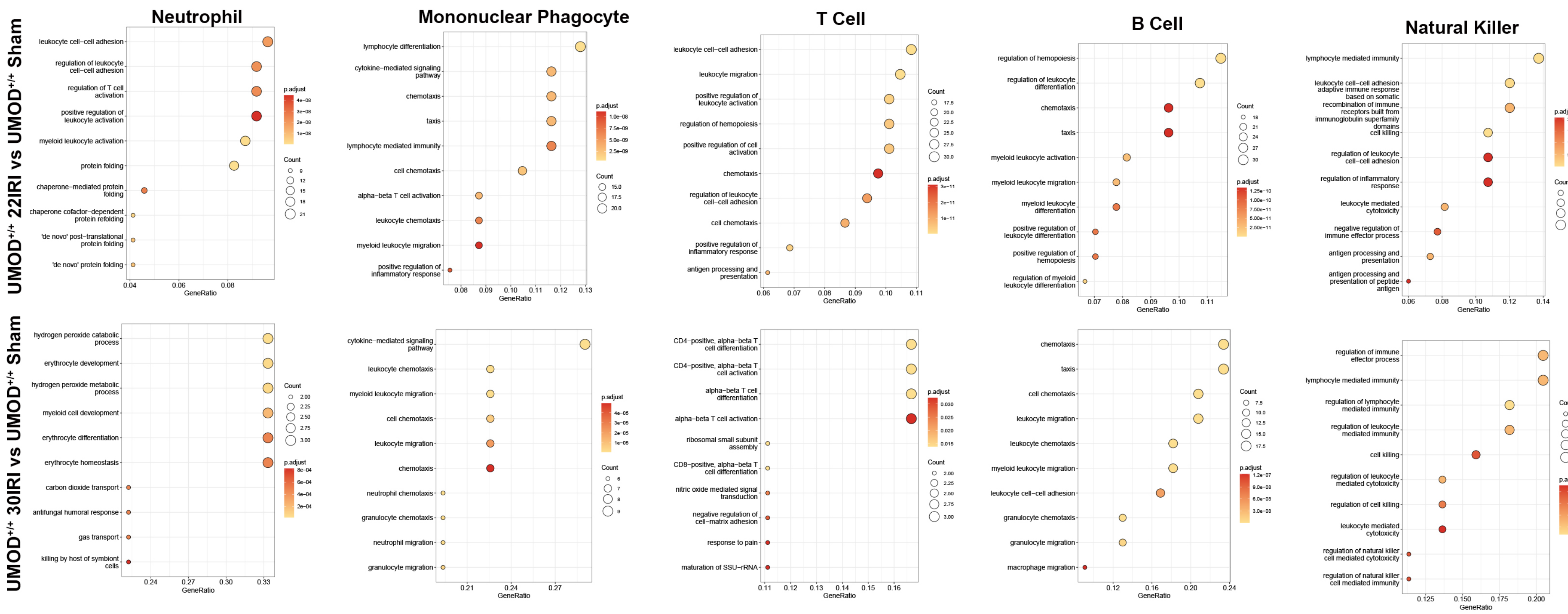


**Fig. S8.** Signaling pathways upregulated in immune cells with moderate (top row) and severe (bottom row) ischemic kidney injury

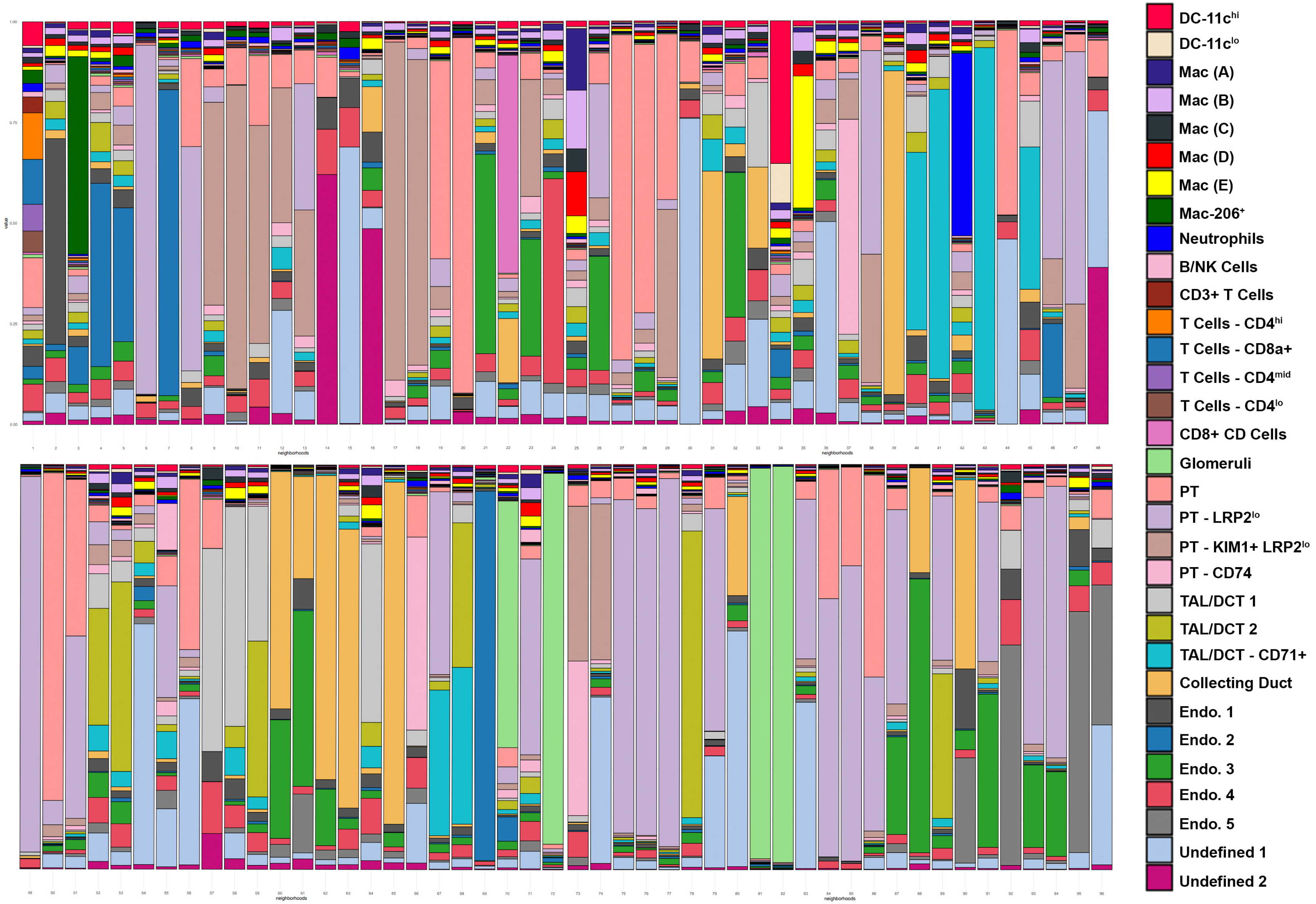


**Fig. S9.** Distribution of cell cluster types within CODEX neighborhoods

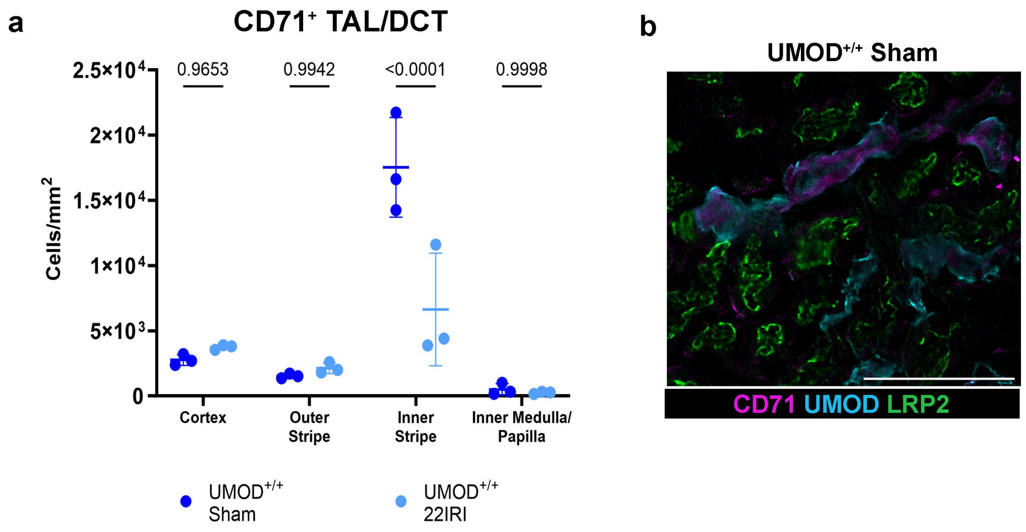


**Fig. S10.** The number of TAL/DCT cells expressing CD71 decreases with injury

- **a)** Distribution of CD71+ TAL/DCT cells decreases in the outer stripe with moderate (22IRI) injury
- **b)** Representative imaging of CD71+ cells in the kidney with markers for TAL/DCT (UMOD) and PT (LRP2)

UMOD – Uromodulin, IRI – ischemia-reperfusion injury, CD – Cluster of Differentiation, TAL – Thick Ascending Limb, DCT – Distal Convoluted Tubule, LRP2 - LDL Receptor Related Protein 2

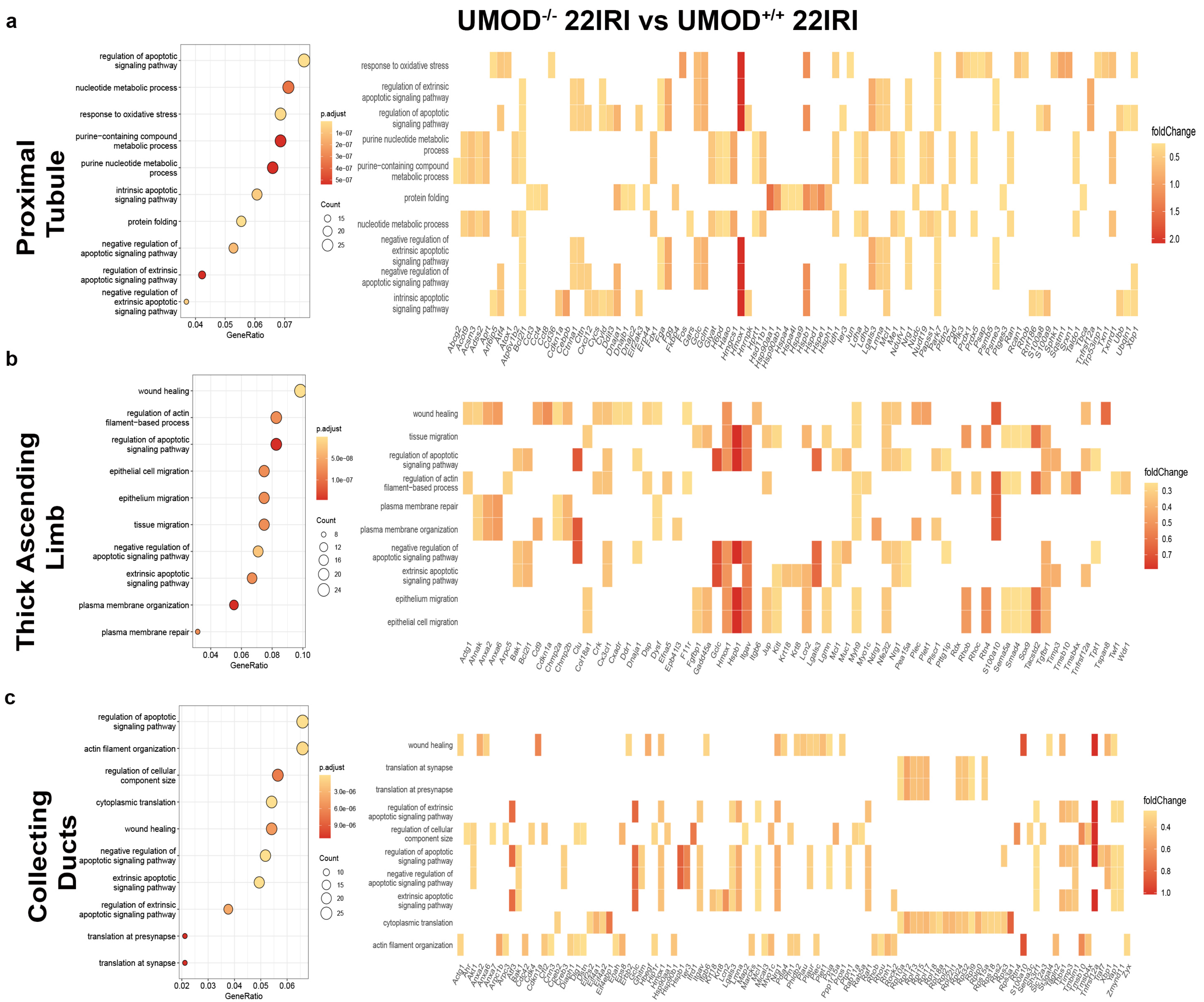

a
b
c
UMOD-/- 22IRI vs UMOD+/+ 22IRI
Proximal Tubule
Thick Ascending Limb
Collecting Ducts
regulation of apoptotic signaling pathway
nucleotide metabolic process
response to oxidative stress
purine−containing compound metabolic process
purine nucleotide metabolic process
intrinsic apoptotic signaling pathway
protein folding
negative regulation of apoptotic signaling pathway
regulation of extrinsic apoptotic signaling pathway
negative regulation of extrinsic apoptotic signaling pathway
GeneRatio
p.adjust
Count
foldChange
wound healing
regulation of actin filament−based process
epithelial cell migration
epithelium migration
tissue migration
extrinsic apoptotic signaling pathway
plasma membrane organization
plasma membrane repair
actin filament organization
regulation of cellular component size
cytoplasmic translation
translation at presynapse
translation at synapse

**Fig. S11.** Uromodulin deficiency increases injury across tubule types

- **a)** Differentially regulated genes contributing to the identification of the top 10 differentially regulated pathways in proximal tubules in injured $UMOD^{-/-}$ mice compared to injured $UMOD^{+/+}$ mice show increased apoptosis, response to oxidative stress and metabolic changes
- **b)** Differentially regulated genes contributing to the identification of the top 10 differentially regulated pathways in thick ascending limb tubules in injured $UMOD^{-/-}$ mice compared to injured $UMOD^{+/+}$ mice show increased apoptosis, wound healing, cell migration and membrane changes
- **c)** Differentially regulated genes contributing to the identification of the top 10 differentially regulated pathways in collecting duct tubules in injured $UMOD^{-/-}$ mice compared to injured $UMOD^{+/+}$ mice show increased apoptosis, wound healing, and cytoskeletal and protein translation changes

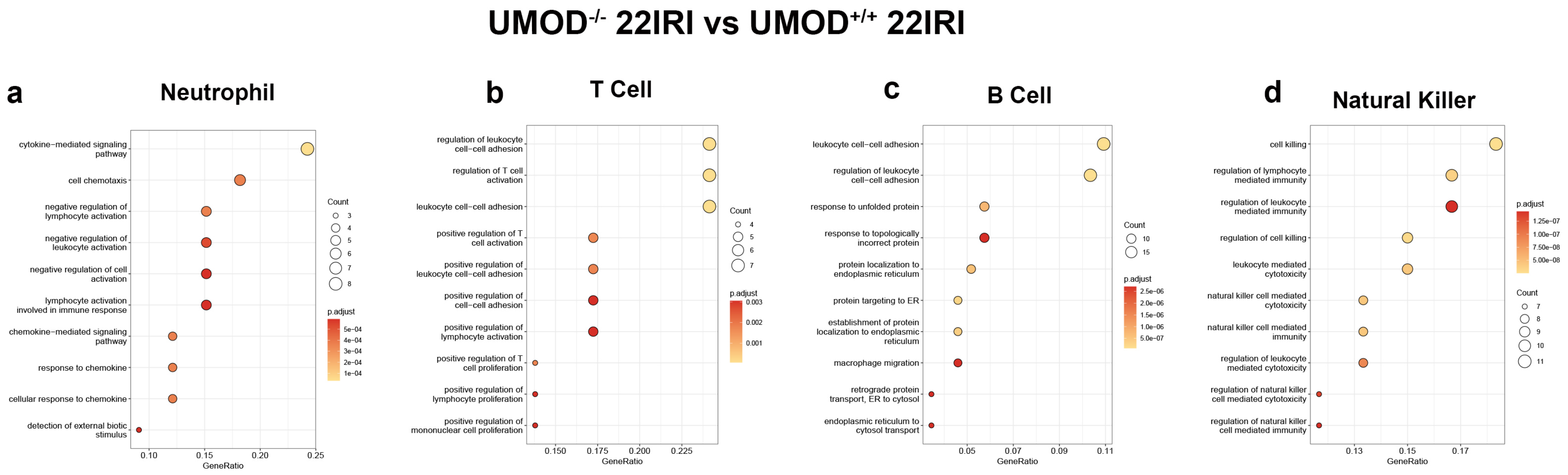


**Fig. S12.** Signaling pathways upregulated in immune cells with Uromodulin deficiency in ischemic kidney injury

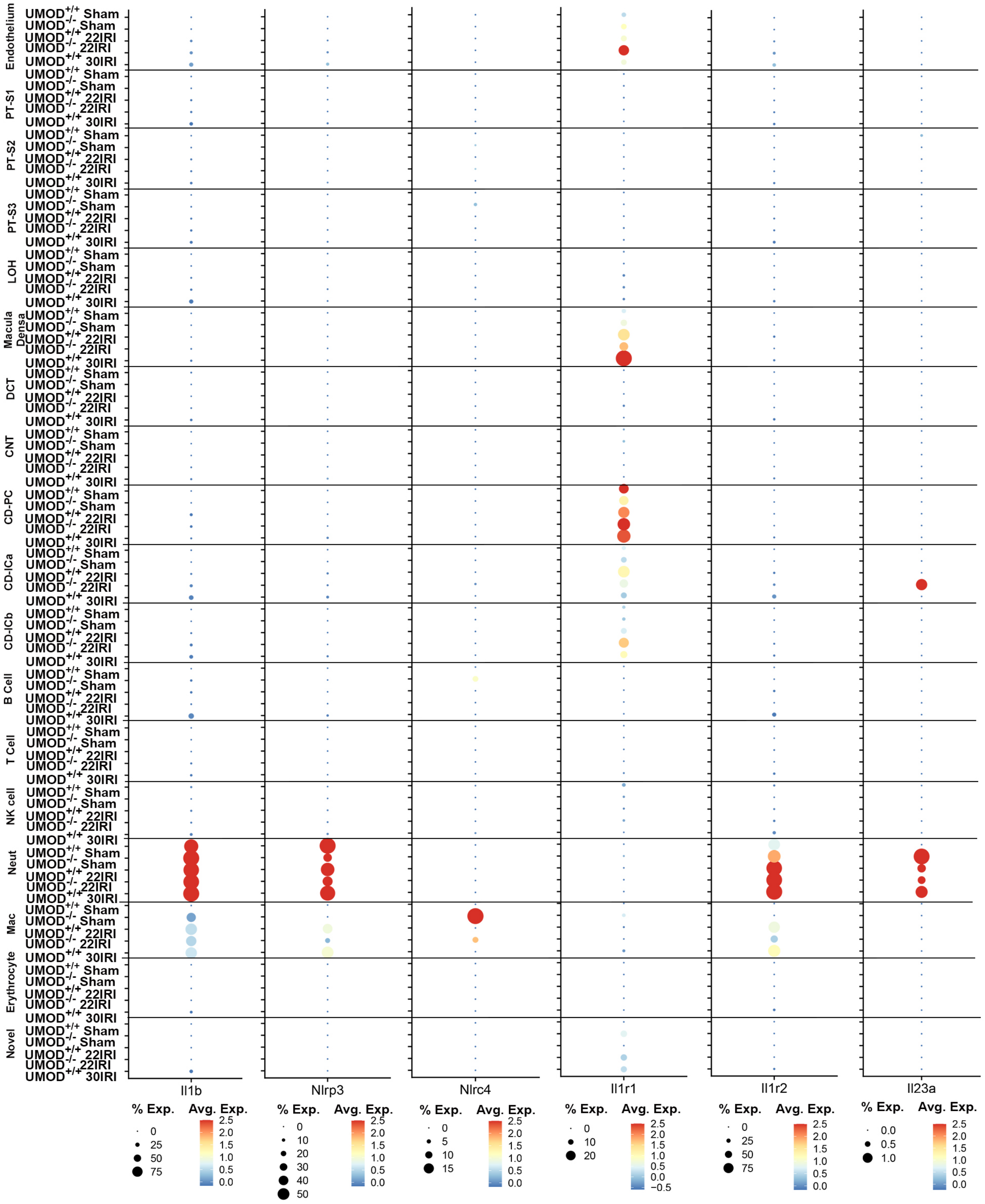


**Fig. S13.** Expression of genes associated with IL1-β release and response across cell types and conditions in whole kidney scRNA-seq

UMOD – Uromodulin, IRI – ischemia-reperfusion injury, PT-S1 – S1 proximal tubule, PT-S2 – S2 proximal tubule, PT-S3 – S3 proximal tubule, LOH – Loop of Henle, DCT – distal convoluted tubule, CNT – connecting tubule, CD-PC – Collecting Duct – Principal Cell, CD-ICa – Collecting Duct – Intercalated Cell Type A, CD-ICb – Collecting Duct – Intercalated Cell Type B

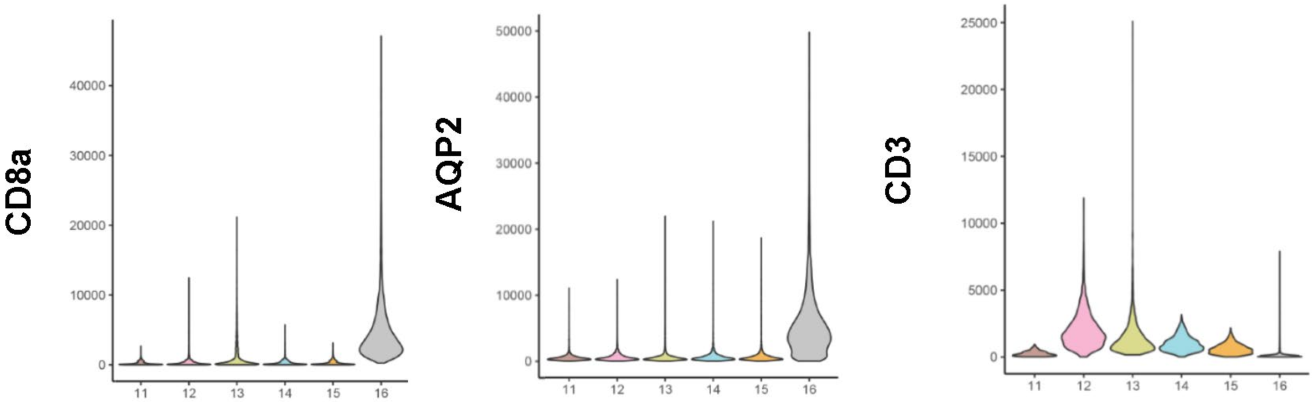


**Fig. S14.** Identification of a CD8a$^{+}$, AQP2$^{+}$, CD3$^{-}$ cell cluster in the kidney by CODEX imaging and cell clustering

CD – Cluster of Differentiation, AQP2 – Aquaporin 2

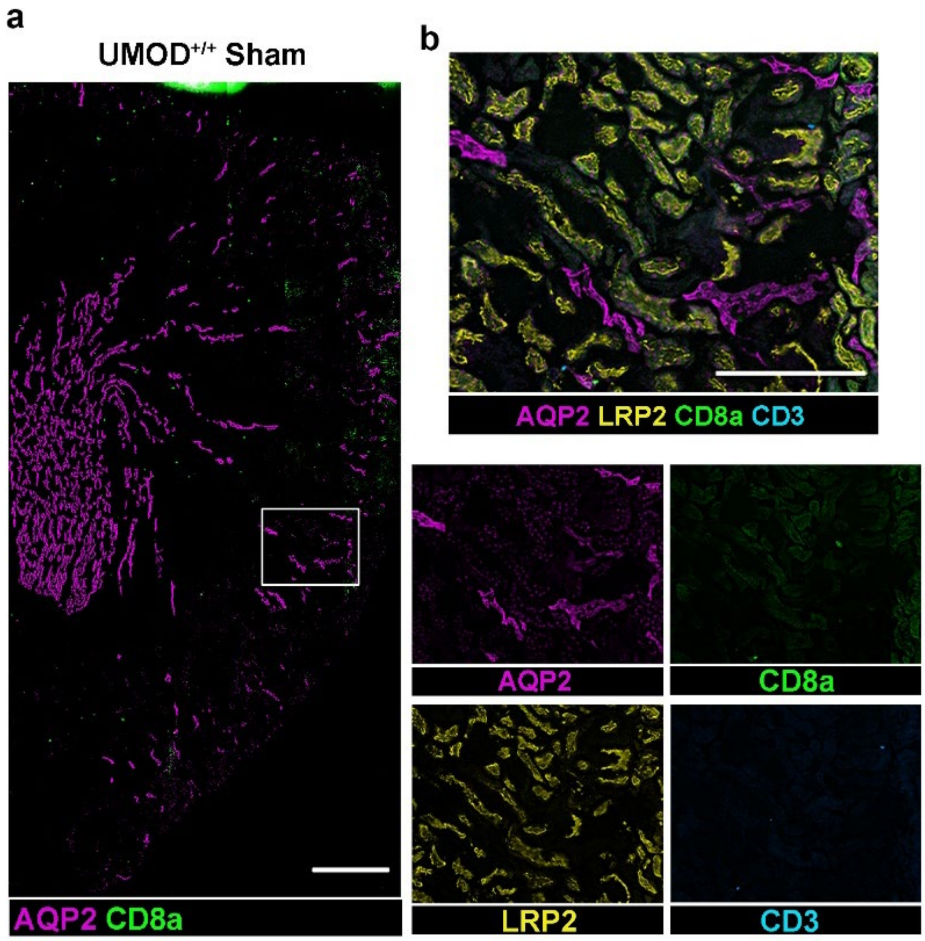


**Fig. S15.** Limited co-localization of AQP2 and CD8a in the kidneys of $UMOD^{+/+}$ Sham-operated mice

- **a)** Representative image of CD8a and AQP2 in $UMOD^{+/+}$ sham kidney
- **b)** Magnification of (a) with additional markers for markers for proximal tubules (LRP2) and T cells (CD3)

UMOD – Uromodulin, AQP2 – aquaporin 2, LRP2 - LDL Receptor Related Protein 2, CD8a – Cluster of Differentiation 8a, CD3 – Cluster of Differentation 3, IRI – Ischemia-reperfusion injury

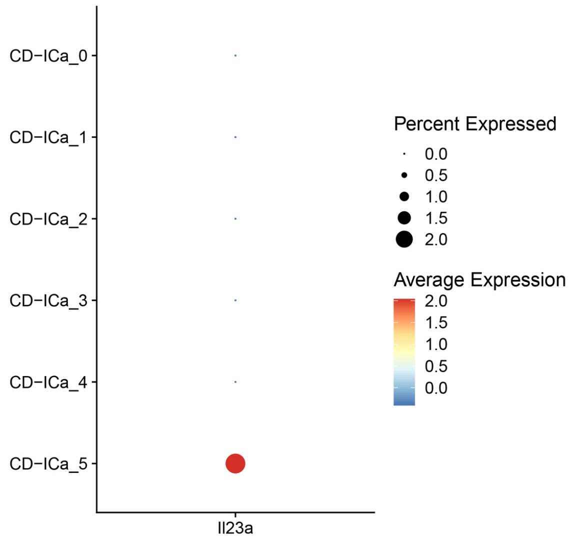


**Fig. S16.** Expression of *Il23a* in Collecting Duct Intercalated Cell Type A subclusters

CD-ICa – Collecting Duct – Intercalated Cell Type A
