## Supplemental File 1 for "Uromodulin promotes immune zonation and inhibits alternative inflammasome-mediated activation of immune-to-collecting duct inflammatory signaling in early acute kidney injury"

MGQPSLTWMLMVVVASWFITTAATDTSEARWCSECHSNATCTEDEAVTTCTCQEGFTGDGLTCVDLDECAIPGAHNCSANSSCVNTPGSFSCVCPEGFRLSPGLGCTDVDECAEPGLSHCHALATCVNVVGSYLCVCPAGYRGDGWHCECSPGSCGPGLDCVPEGDALVCADPCQAHRTLDEYWRSTEYGEGYACDTDLRGWYRFVGQGGARMAETCVPVLRCNTAAPMWLNGTHPSSDEGIVSRKACAHWSGHCCLWDASVQVKACAGGYYVYNLTAPPECHLAYCTDPSSVEGTCEECSIDEDCKSNNGRWHCQCKQDFNITDISLLEHRLECGANDMKVSLGKCQLKSLGFDKVFMYLSDSRCSGFNDRDNRDWVSVVTPARDGPCGTVLTRNETHATYSNTLYLADEIIIRDLNIKINFACSYPLDMKVSHHHHHH
