## Supplemental File 2 for "Uromodulin promotes immune zonation and inhibits alternative inflammasome-mediated activation of immune-to-collecting duct inflammatory signaling in early acute kidney injury"

| Gene Target (Mouse) | Manufacturer | Primer ID |
| --- | --- | --- |
| *Bak1* | ThermoFisher Scientific | Mm00432045_m1 |
| *Bax* | ThermoFisher Scientific | Mm00432051_m1 |
| *Il1b* | ThermoFisher Scientific | Mm00434228_m1 |
| *Il23a* | ThermoFisher Scientific | Mm00518984_m1 |
| *Mlkl* | ThermoFisher Scientific | Mm01244222_m1 |
| *Nlrp3* | ThermoFisher Scientific | Mm00840904_m1 |
| *Nlrc4* | ThermoFisher Scientific | Mm01239561_m1 |
| *Ripk1* | ThermoFisher Scientific | Mm00436354_m1 |
